# Conserved properties of *Drosophila* Insomniac link sleep regulation and synaptic function

**DOI:** 10.1101/104893

**Authors:** Qiuling Li, David A. Kellner, Hayden A. M. Hatch, Tomohiro Yumita, Sandrine Sanchez, Robert P. Machold, C. Andrew Frank, Nicholas Stavropoulos

## Abstract

Sleep is an ancient animal behavior that is regulated similarly in species ranging from flies to mammals. Various genes that regulate sleep have been identified in invertebrates, but whether the functions of these genes are conserved in mammals remains poorly explored. *Drosophila insomniac (inc)* mutants exhibit severely shortened and fragmented sleep. Inc protein physically associates with the Cullin-3 (Cul3) ubiquitin ligase, and neuronal depletion of Inc or Cul3 strongly curtails sleep, suggesting that Inc is a Cul3 adaptor that directs the ubiquitination of neuronal substrates that regulate sleep. Three proteins similar to Inc exist in vertebrates—KCTD2, KCTD5, and KCTD17—but are uncharacterized within the nervous system and their functional conservation with Inc has not been addressed. Here we show that Inc and its mouse orthologs exhibit striking biochemical and functional interchangeability within Cul3 complexes. Remarkably, KCTD2 and KCTD5 restore sleep to *inc* mutants, indicating that they can substitute for Inc in vivo and engage its neuronal targets that impact sleep. Inc and its orthologs traffic similarly within fly and mammalian neurons and are present at synapses, suggesting that their substrates include synaptic proteins. Consistent with such a mechanism, *inc* mutants exhibit defects in synaptic structure and physiology, indicating that Inc is vital for both synaptic function and sleep. Our findings reveal that molecular functions of Inc are conserved through ~600 million years of evolution and support the hypothesis that Inc and its orthologs participate in an evolutionarily conserved ubiquitination pathway that links synaptic function and sleep regulation.

**Author summary:** Sleep is ubiquitous among animals and is regulated in a similar manner across phylogeny, but whether conserved molecular mechanisms govern sleep is poorly defined. The Insomniac protein is vital for sleep in *Drosophila* and is a putative adaptor for the Cul3 ubiquitin ligase. We show that two mammalian orthologs of Insomniac can restore sleep to flies lacking Insomniac, indicating that the molecular functions of these proteins are conserved through evolution. Our comparative analysis reveals that Insomniac and its mammalian orthologs localize to neuronal synapses and that Insomniac impacts synaptic structure and physiology. Our findings suggest that Insomniac and its mammalian orthologs are components of an evolutionarily conserved ubiquitination pathway that links synaptic function and the regulation of sleep.

## Introduction

Sleep is an evolutionarily ancient behavior present in vertebrates and invertebrates [1]. The similar characteristics of sleep states across animal phylogeny suggest that both the functions of sleep and the regulation of sleep may have a common evolutionary basis [2]. In diverse animals including mammals and insects, sleep is regulated similarly by circadian and homeostatic mechanisms [3]. The circadian regulation of sleep is better understood at a molecular level, and numerous studies have revealed that the underlying genes and intracellular pathways are largely conserved from flies to humans [4-10]. In contrast, the molecular mechanisms underlying the non-circadian regulation of sleep—including those governing sleep duration, consolidation, and homeostasis—remain less well defined, and their evolutionary conservation is largely unexplored.

Mutations of the *Drosophila insomniac* (*inc*) gene [11] severely curtail the duration and consolidation of sleep but do not alter its circadian regulation [11,12]. *inc* encodes a protein of the Bric-à-brac, Tramtrack, and Broad / Pox virus zinc finger (BTB/POZ) superfamily [13], which includes adaptors for the Cullin-3 (Cul3) E3 ubiquitin ligase complex [14-17]. Cul3 adaptors have a modular structure, in which the BTB domain binds Cul3 and a second distal domain recruits substrates to the Cul3 complex for ubiquitination [14-17]. The BTB domain also mediates adaptor self-association, enabling the oligomerization of Cul3 complexes and the efficient recruitment and ubiquitination of substrates [18,19]. Biochemical and genetic evidence supports the hypothesis that Inc is a Cul3 adaptor [11,12]. Inc and Cul3 physically interact in cultured cells [11,12] and Inc is able to self-associate [12]. In vivo, neuronal RNAi against *inc* or *Cul3* strongly reduces sleep [11,12], and reduction in the levels of Nedd8, a protein whose conjugation to Cullins is essential for their activity, also decreases sleep [11]. While Inc is thus likely to function as a Cul3 adaptor within neurons to promote sleep, the neuronal mechanisms through which Inc influences sleep are unknown.

Three proteins similar to Inc—KCTD2, KCTD5, and KCTD17—are present in vertebrates [11,20,21], but their functions in the nervous system are uncharacterized and their functional conservation with Inc has not been addressed. KCTD5 can self-associate and bind Cul3 [20], suggesting that it may serve as a Cul3 adaptor, yet no substrates have been identified among its interacting partners [22,23]. One KCTD17 isoform has been shown to function as a Cul3 adaptor for trichoplein, a regulator of primary cilia [24]. However, trichoplein-binding sequences are not present in Inc, KCTD2, KCTD5, or other KCTD17 isoforms, and trichoplein is not conserved in *Drosophila*. Thus, it remains unclear whether Inc and its vertebrate homologs have conserved molecular functions, particularly within neurons and cellular pathways relevant to sleep.

Here, we assess the functional conservation of Inc and its mammalian orthologs and elucidate a neuronal mechanism through which they may impact sleep. Inc and each of its orthologs bind Cul3 and self-associate, supporting a universal role for these proteins as Cul3 adaptors. Inc and its orthologs furthermore exhibit biochemical interchangeability within Inc-Inc and Inc-Cul3 complexes, indicating that the oligomeric architecture of Inc-Cul3 complexes is highly conserved. Strikingly, KCTD2 and KCTD5 can functionally substitute for Inc in vivo and restore sleep to *inc* mutants, indicating that these Inc orthologs readily engage the molecular targets through which Inc impacts sleep. Our studies furthermore reveal that Inc and its orthologs localize similarly within fly and mammalian neurons and traffic to synapses. Finally, we show that *inc* mutants exhibit defects in synaptic structure and physiology, indicating that *inc* is essential for both sleep and synaptic function. Our findings demonstrate that molecular functions of Inc are conserved from flies to mammals, and support the hypothesis that Inc and its orthologs direct the ubiquitination of conserved neuronal proteins that link sleep regulation and synaptic function.

## Results

### The mammalian orthologs of Insomniac are expressed in the nervous system

Inc functions in neurons to impact sleep [11, 12]. To assess whether Inc orthologs are expressed in the mammalian nervous system, we performed RT-PCR on mouse brain RNA using primers specific for KCTD2, KCTD5, and KCTD17. All three genes are expressed in the brain (Fig 1A), and in situ hybridizations reveal expression within cortex, thalamus, striatum, pons, and cerebellum among other brain regions (S1 Fig). Cloning of RT-PCR products revealed single transcripts for KCTD2 and KCTD5, encoding proteins of 263 and 234 residues respectively, and two alternatively spliced transcripts for KCTD17, encoding proteins of 225 and 220 residues with distinct C-termini (Fig 1B and S2A Fig). These KCTD17 isoforms, KCTD17.2 and KCTD17.3, have not been characterized previously and lack residues of a longer KCTD17 isoform required to bind trichoplein (S2B Fig) [24]. Thus, KCTD17.2 and KCTD17.3 are likely to have different molecular partners. KCTD17.2 and KCTD17.3 behaved indistinguishably in our experiments except where noted below. Inc and its mouse orthologs share ~60% sequence identity, and have variable N-termini followed by a BTB domain, an intervening linker, and conserved C-termini (S2A Fig) [11,20,21].

**Fig 1.**
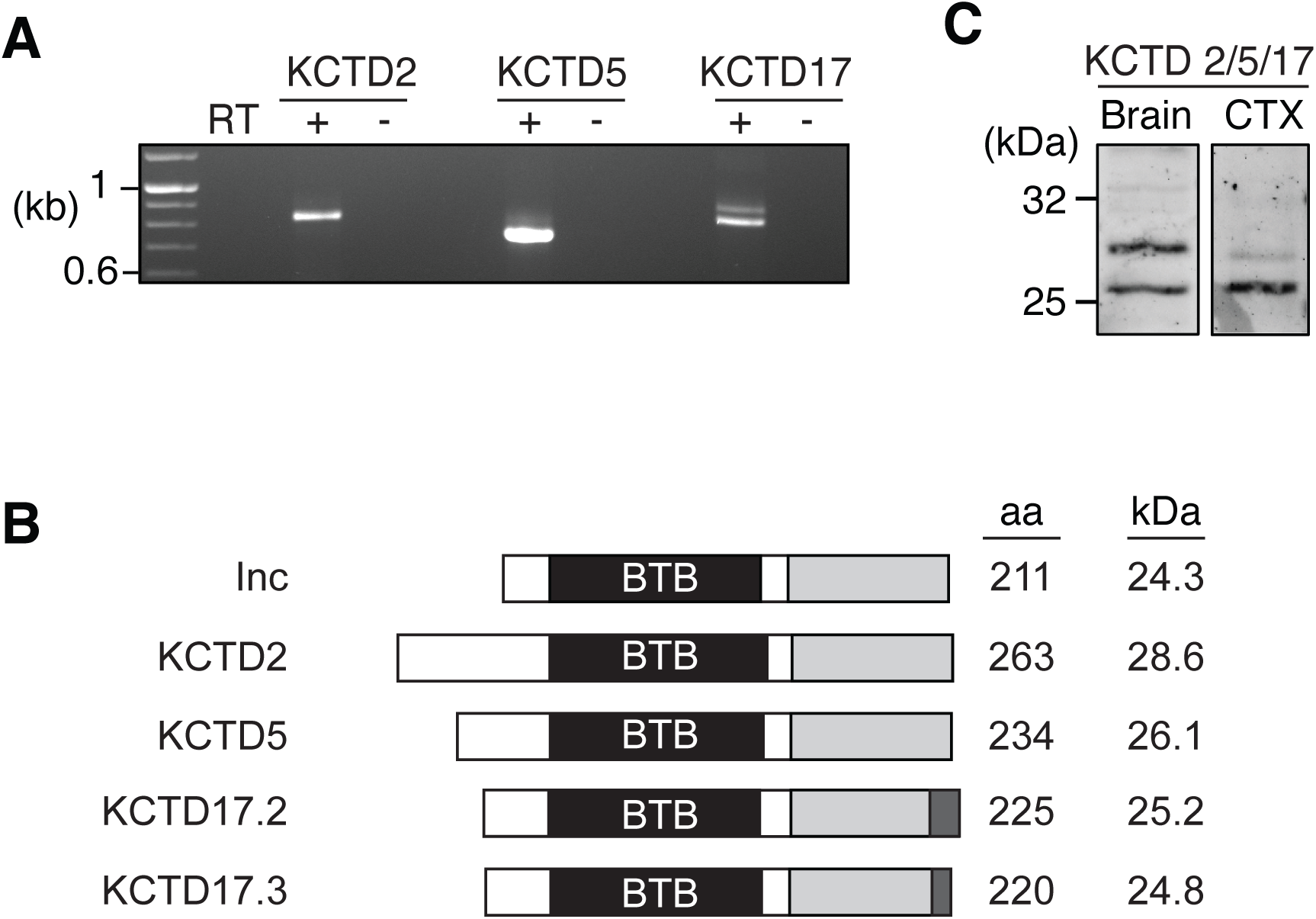
Expression analysis of mammalian Insomniac orthologs. (**A**) RT-PCR amplification of KCTD2, KCTD5, and KCTD17 from total mouse brain RNA. The presence (+) or absence (-) of reverse transcriptase (RT) prior to PCR is indicated. (**B**) Structure of Inc family members. Conserved BTB (black) and C-terminal domains (grey), KCTD17 alternative variant residues (dark grey), number of amino acid (aa) residues, and predicted molecular weights are indicated. (**C**) Western blot of mouse brain and rat cortical culture (CTX) extracts probed with anti-KCTD5 antibody that recognizes KCTD2, KCTD5, and KCTD17.

We next assessed the expression of KCTD2, KCTD5, and KCTD17 proteins, using a polyclonal anti-KCTD5 antibody that cross-reacts with mouse Inc orthologs and *Drosophila* Inc (S3A Fig). This antibody detected a strongly reactive species of ~26 kD and additional species of ~28 to 29 kD in extracts from mouse and rat brain, cultured rat cortical neurons, and human 293T cells (Figs 1C and 5D, and S3A Fig), consistent with the range of molecular weights predicted for KCTD2, KCTD5, and KCTD17 (Fig 1B). The size of these immunoreactive species and their biochemical properties, described further below, indicate that they correspond to one or more isoforms of KCTD2, KCTD5, and KCTD17. The expression of Inc orthologs in the mammalian brain and Inc in the fly brain [11], together with the similarity of their primary sequences, suggests that Inc defines a protein family that may have conserved functions in the nervous system.

### Insomniac family members self-associate and bind Cul3 in an evolutionarily conserved and interchangeable manner

Inc and KCTD5 are able to bind Cul3 and to self-associate [11, 12, 20], key attributes of BTB adaptors [14-17]. To determine whether these attributes are universal to Inc orthologs and their isoforms expressed in the nervous system, we first examined the physical interactions of these proteins with mouse Cul3. Co-immunoprecipitations revealed that KCTD2, KCTD5, KCTD17.2, and KCTD17.3 are able to associate with mouse Cul3 (Fig 2A and data not shown), with KCTD5 exhibiting stronger or more stable interactions than KCTD2 and KCTD17. Thus, the ability to bind Cul3 is a conserved property of Inc and its mouse orthologs. Epitope-tagged Cul3 also co-immunoprecipitated the endogenous ~26 kD species detected by anti-KCTD5 antibody, confirming that this species represents KCTD5 or another Inc ortholog (S3B Fig).

**Fig 2.**
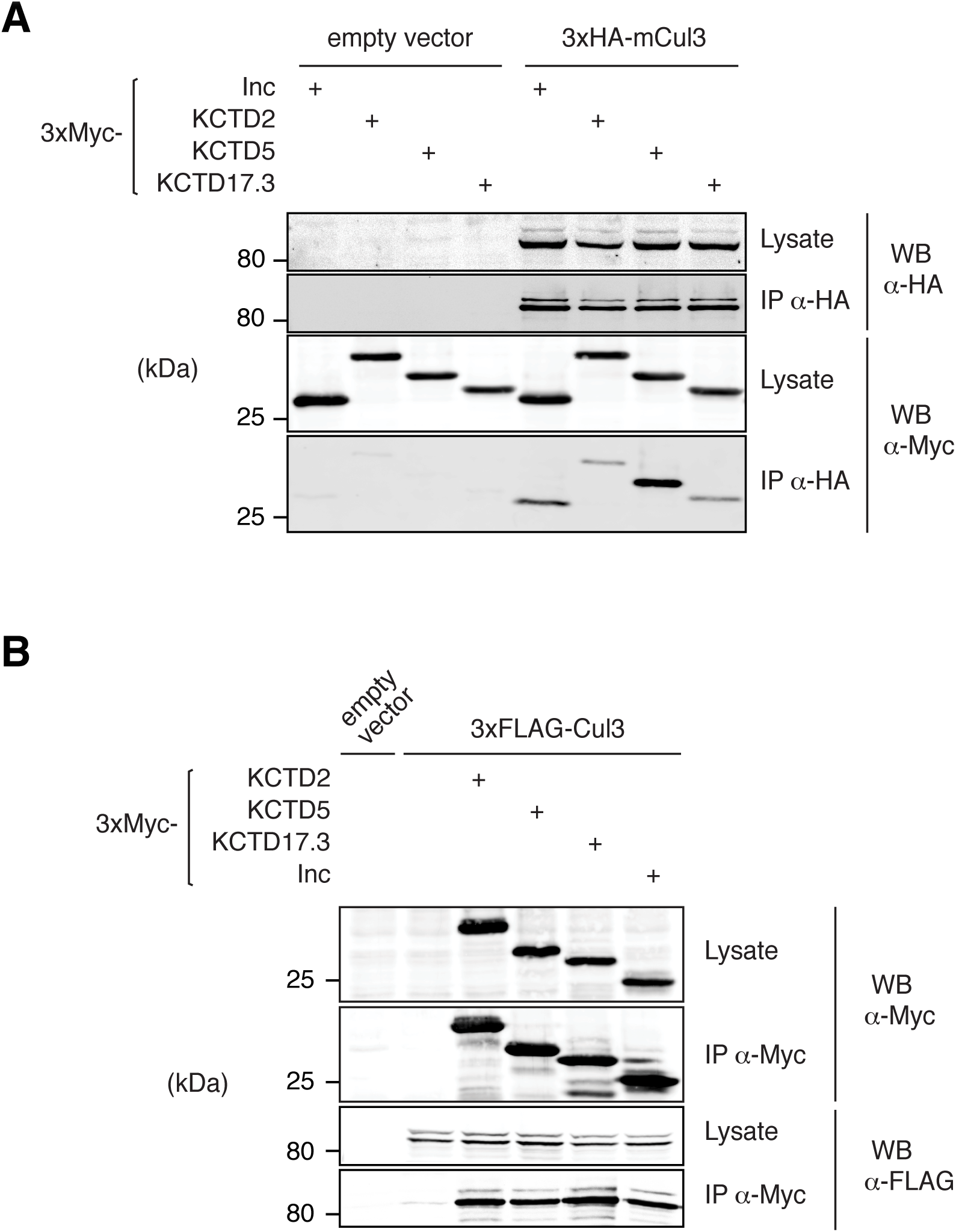
Insomniac-Cul3 interactions are conserved in flies and mammals. (**A and B**) Co-immunoprecipitation analysis of epitope-tagged Inc or mouse Inc orthologs co-expressed (A) with mouse Cul3 (mCul3) in 293T cells or (B) with *Drosophila* Cul3 in Schneider S2 cells.

To assess the extent to which the Inc-Cul3 interface is evolutionarily conserved, we tested whether *Drosophila* Inc is able to associate with mouse Cul3 in a cross-species manner. We observed that fly Inc and mouse Cul3 interact (Fig 2A), indicating that Inc readily assembles into mammalian Cul3 complexes. Conversely, we tested whether mouse KCTD2, KCTD5, and KCTD17 can associate with fly Cul3 in *Drosophila* S2 cells, and observed that each Inc ortholog associated with fly Cul3 in a manner indistinguishable from Inc (Fig 2B). The interchangeable biochemical associations of Inc family members and Cul3 indicate that the Inc-Cul3 interface is functionally conserved from flies to mammals.

Self-association is a critical property of BTB adaptors that enables the oligomerization of Cul3 complexes and that stimulates substrate ubiquitination [18, 19]. To test whether Inc orthologs self-associate in a manner similar to Inc [12], we co-expressed FLAG-and Myc-tagged forms of each Inc ortholog in mammalian cells and performed co-immunoprecipitations. KCTD2, KCTD5, and KCTD17 each homomultimerized strongly (Fig 3A). Thus, homo-oligomerization is a shared property of Inc family proteins. The presence of three Inc orthologs in mammals and their likely co-expression in brain regions such as thalamus and cortex (S1 Fig) led us to test whether these proteins can also heteromultimerize. We observed robust heteromeric associations between all pairwise combinations of Inc orthologs (Fig 3A), a property that may enable functional redundancy in vivo or the assembly of functionally distinct complexes. To further probe the multimeric self-associations of Inc family members, we tested whether KCTD2, KCTD5, and KCTD17 can heteromultimerize with *Drosophila* Inc. We observed that each Inc ortholog associates readily with Inc in both mammalian and *Drosophila* cells (Fig 3A and 3B). Thus, the multimerization interface of Inc family members is highly conserved through evolution. Together with the interchangeable associations of Inc family members and Cul3 (Fig 2), these findings strongly suggest a conserved oligomeric architecture for complexes containing Cul3 and Inc family members.

**Fig 3.**
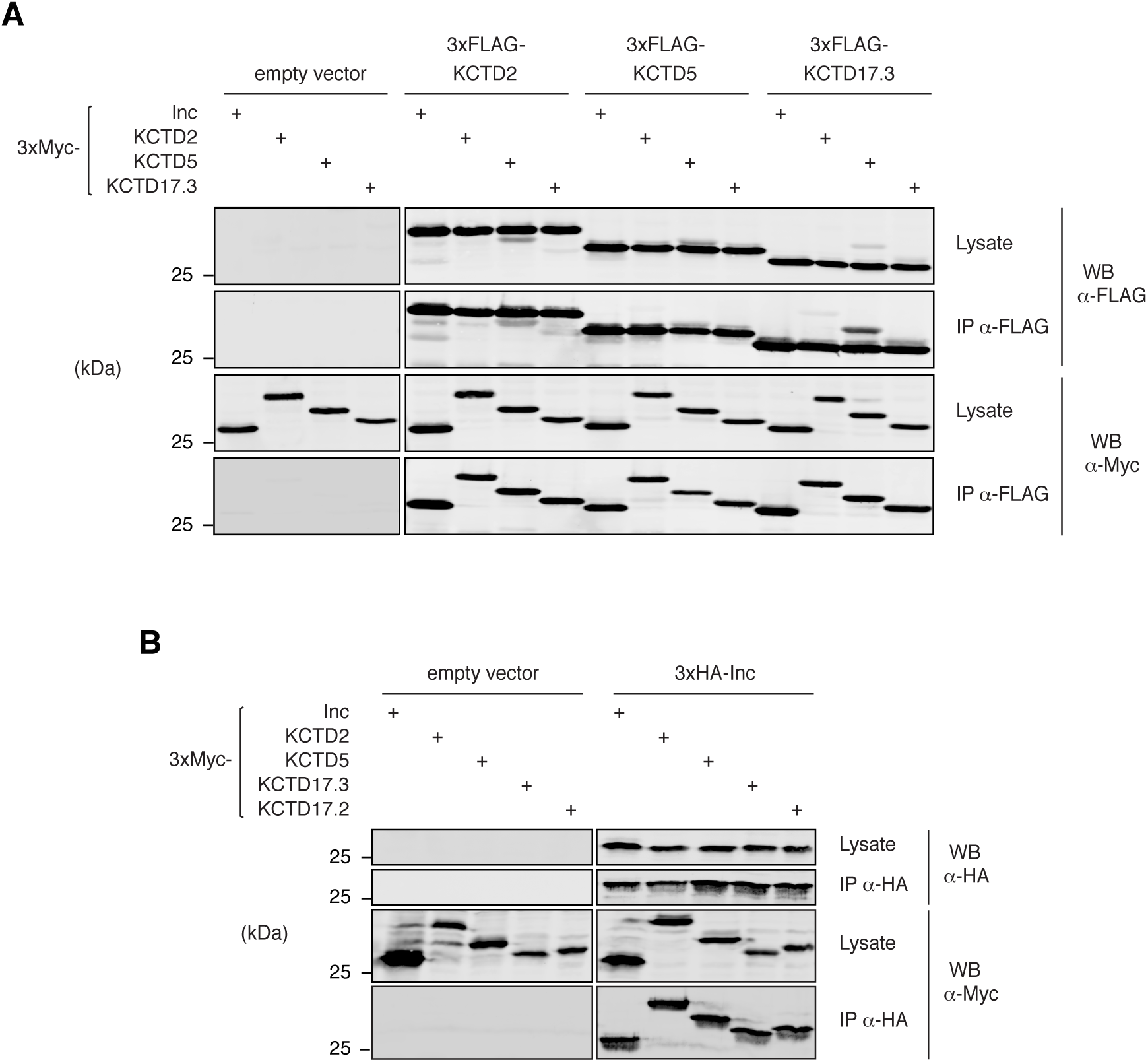
Homomeric and heteromeric interactions of Insomniac family members. (**A and B**) Co-immunoprecipitation analysis of epitope-tagged Inc and mouse Inc orthologs co-expressed in a pairwise manner in (A) 293T cells or (B) *Drosophila* S2 cells.

### KCTD2 and KCTD5 substitute for *Drosophila* Inc in vivo and restore sleep to *inc* mutants

The biochemical interchangeability of Inc and its mouse orthologs prompted us to test whether Inc function, including its presumed ability to serve as a Cul3 adaptor in vivo and ubiquitinate specific neuronal proteins relevant to sleep, is conserved between flies and mammals. We therefore generated UAS transgenes expressing Myc-tagged forms of Inc, KCTD2, KCTD5, KCTD17.2, and KCTD17.3, integrated each at the *attP2* site [25], and backcrossed these lines to generate an isogenic allelic series. Expression of these transgenes panneuronally with *elav*^*c155*^*-Gal4* yielded similar levels of expression for Inc, KCTD2, and KCTD5, weak expression of KCTD17.2, and low levels of KCTD17.3 expression near the threshold of detection (Fig 4A). Next, we assessed the behavioral consequences of expressing mouse Inc orthologs in vivo. Animals expressing Inc orthologs under the control of *elav*^*c155*^*-Gal4* slept largely indistinguishably from control animals expressing Inc or from those lacking a UAS transgene, as indicated by analysis of sleep duration, daytime and nighttime sleep, sleep bout length, and sleep bout number (Fig 4C and S4A-S4D Fig). Thus, neuronal expression of Inc and its mouse orthologs does not elicit significant dominant negative effects or otherwise inhibit endogenous *inc* function, similar to ubiquitous or neuronal expression of untagged Inc [11].

**Fig 4.**
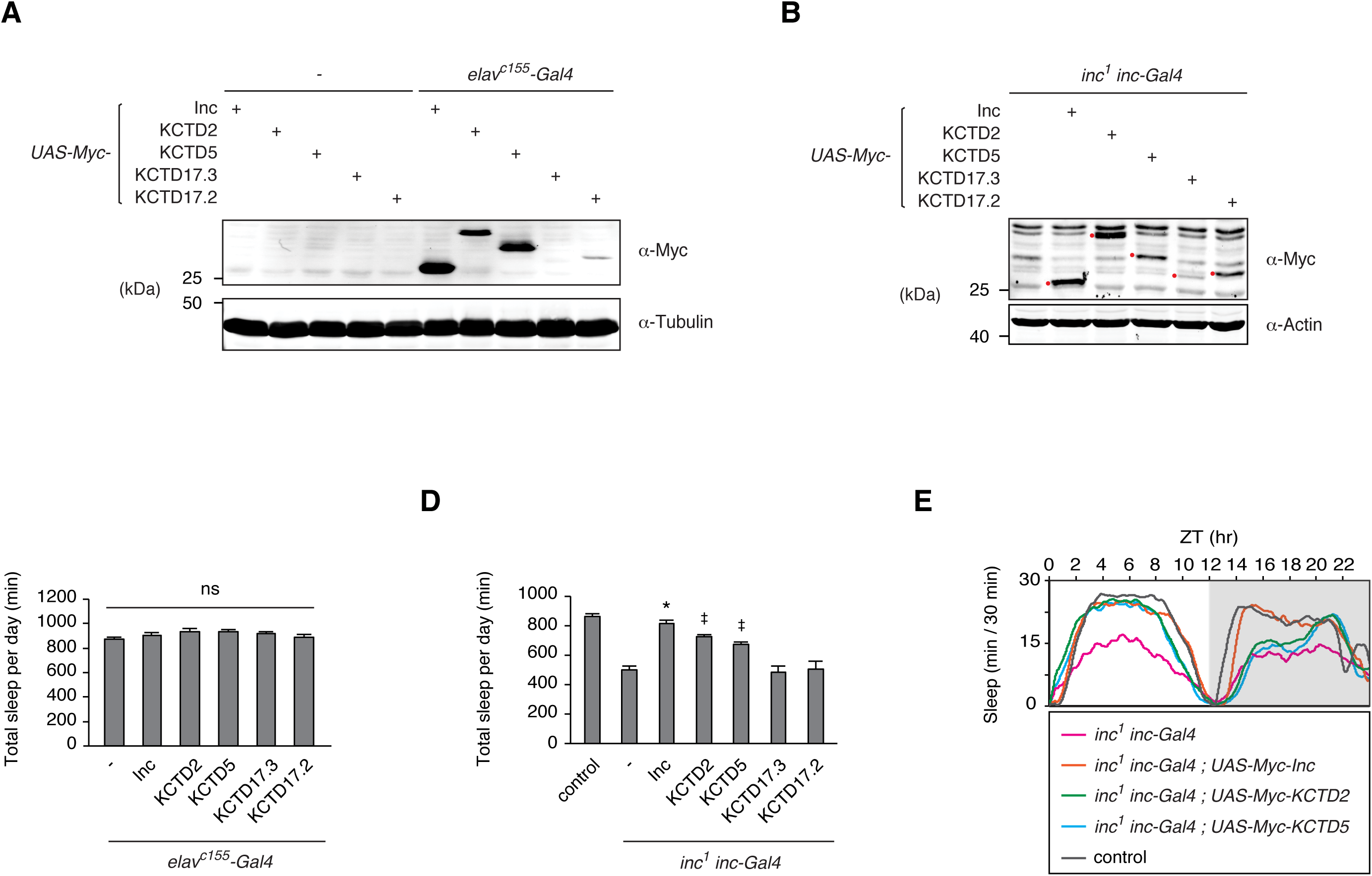
Insomniac orthologs functionally substitute for Insomniac and restore sleep to inc mutants in vivo. (**A and B**) Western blots of head (A) or whole animal lysates (B) prepared from indicated genotypes.Myc-tagged Inc family members are indicated with red dots in (B). (**C and D**) Total sleep per day for indicated genotypes. Mean ± SEM is shown. For (C), n = 37-40; ns, not significant (p > 0.05). For (D) n = 18-157; * p < 0.01 for comparison to *inc*^*1*^*inc-Gal4* animals and not significantly different from wild-type controls; ^‡^ p < 0.01 for comparisons to *inc*^*1*^ *inc-Gal4* animals and to wild-type controls. (E) Population average sleep traces for indicated genotypes. n = 56-157.

To assess whether mouse Inc orthologs can functionally substitute for *Drosophila* Inc, we measured their ability to rescue the sleep defects of *inc*^*1*^ null mutants [11]. Sleep is a behavior sensitive to genetic background [26,27], environment [28], and other influences, and thus the ability of Inc orthologs to confer behavioral rescue represents a stringent test of *inc* function. Inc and its orthologs expressed under *inc-Gal4* control in *inc*^*1*^ animals accumulated with relative levels similar to those in *elav-Gal4* animals (Fig 4A and 4B). Expression of Myc-Inc under *inc-Gal4* control fully rescued sleep in *inc* mutants to wild-type levels, indicating that Myc-Inc recapitulates the function of endogenous Inc protein (Fig 4D). Strikingly, expression of mouse KCTD2 and KCTD5 strongly rescued *inc* sleep defects, including total sleep duration (Fig 4D), the distribution of sleep throughout the day (Fig 4E), and most other sleep parameters (S4E-S4H Fig). This rescue indicates that KCTD2 and KCTD5 not only recapitulate Inc-Inc and Inc-Cul3 interactions (Figs 2 and 3), but that they retain other critical aspects of Inc function including its presumed ability to engage and ubiquitinate neuronal target proteins in vivo. In contrast, KCTD17.2 and KCTD17.3 failed to rescue *inc* phenotypes (Fig 4D and S4E-S4H Fig). The restoration of sleep-wake cycles by KCTD2 and KCTD5 but not by isoforms of KCTD17 contrasts with the apparent biochemical interchangeability of these proteins with respect to Inc-Inc and Inc-Cul3 associations (Figs 2 and 3). The inability of KCTD17.2 and KCTD17.3 to rescue *inc* sleep defects may reflect the lower abundance of these proteins in transgenic flies (Fig 4A and 4B) or differences in the intrinsic activities of these KCTD17 isoforms, including their ability to engage Inc targets relevant to sleep.

### Insomniac family members traffic through neuronal projections and localize to synapses in mammalian neurons

Inc is required in neurons for normal sleep-wake cycles [11, 12], indicating that it impacts aspects of neuronal function that are essential for sleep. The subcellular localization of Cul3 adaptors reflects their biological functions and variously includes the cytoplasm [29], nuclear foci [30], cytoskeletal structures [31,32], and perisynaptic puncta in neurons [33]. The subcellular distribution of Inc is unknown, as available antisera do not efficiently detect endogenous Inc [11], and the localization of Inc orthologs in neurons is similarly uncharacterized. Anti-KCTD5 antibody did not efficiently detect endogenously expressed Inc orthologs in immunohistochemical staining (data not shown). We therefore examined the subcellular localization of Inc family members fused to epitope tags and expressed in cultured cells, primary neurons, and in flies in vivo. In S2 cells, Myc-tagged Inc was localized to the cytoplasm and excluded from the nucleus (S5A Fig), and an Inc-GFP fusion was distributed similarly in both live cells and after fixation (S5A Fig). Inc orthologs, Inc-GFP, and mouse Cul3 were similarly distributed in the cytoplasm and excluded from the nucleus in cultured mammalian cells (S5B-S5D Fig).

We next assessed the localization of tagged Inc orthologs in primary neurons cultured from cortex, a region of the brain in which they are expressed in vivo (Fig 1C and S1 Fig). In cortical neurons, KCTD2, KCTD5, and KCTD17 were excluded from the nucleus and localized to the cytosol and to dendritic and axonal projections (Fig 5A). Within dendrites, these proteins trafficked to spines as indicated by co-staining against PSD95, a component of the postsynaptic density (Fig 5B). In axons, Inc orthologs were enriched at varicosities costaining with synapsin, a vesicle-associated protein that marks presynaptic termini (Fig 5C). Inc localized similarly to its mammalian orthologs including at pre-and post-synaptic structures (Fig 5A-5C), suggesting that intrinsic determinants governing the localization of Inc and its orthologs in neurons may be functionally conserved.

**Fig 5.**
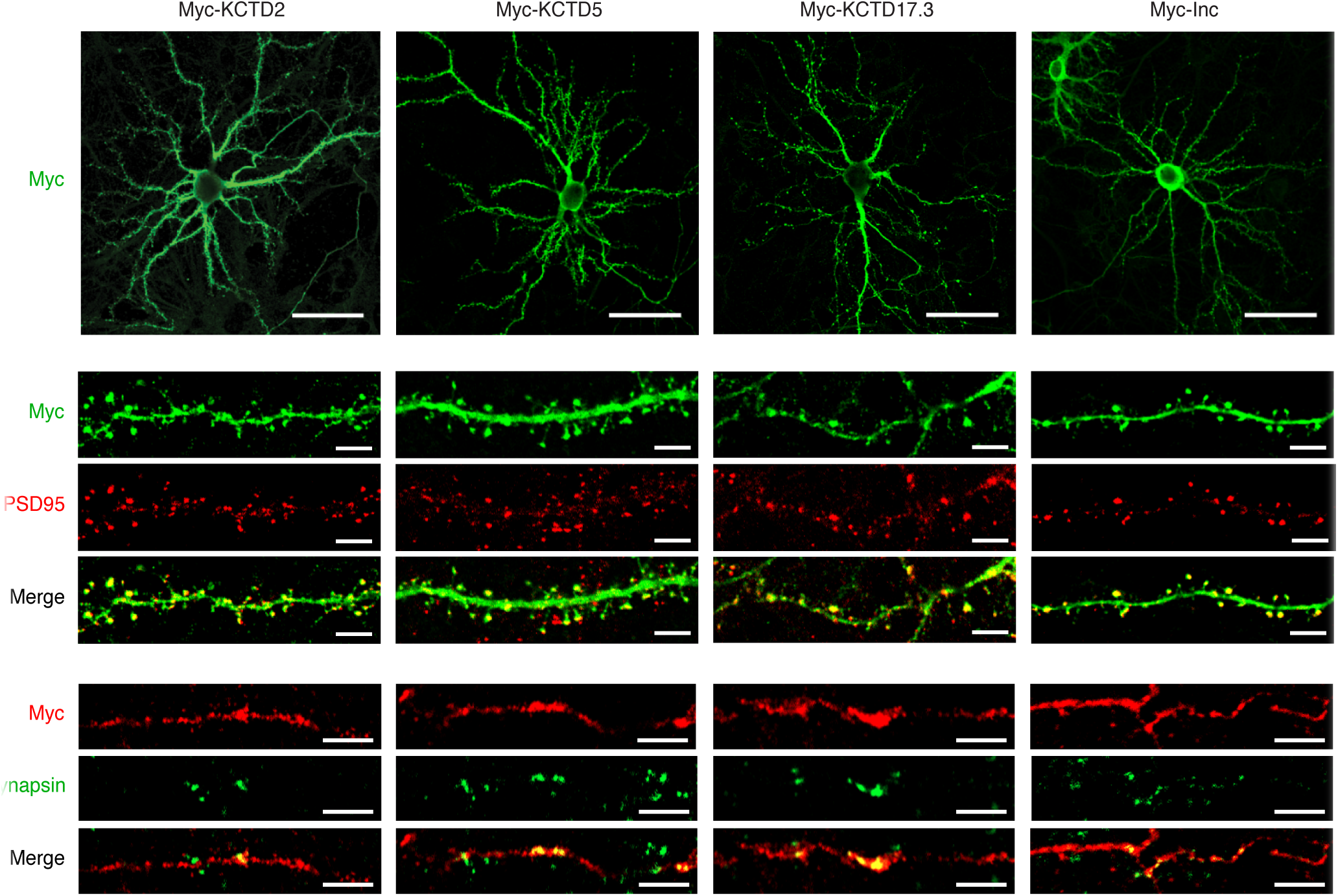
Mammalian Insomniac orthologs traffic to neuronal arborizations and synapses. (**A-C**) Immunohistochemical analysis of rat cortical neurons expressing indicated Myc-tagged proteins. Widefield images of transfected cortical neurons (A), and magnifications of dendrites (B) and axons (C). Scale bars are 50 µm in (A) and 5 µm in (B and C). (**D**) Western blot of total membrane (TM) or synaptosome (SNAP) fractions prepared from rat brain. Note the presence of higher-molecular weight neddylated Cul3.

To assess whether the synaptic localization of Inc orthologs in transfected neurons reflects the distribution of corresponding endogenous proteins, we fractionated cortex to isolate synaptosomes and subjected them to biochemical analysis. Enrichment of presynaptic and postsynaptic proteins in these preparations was confirmed by blotting for synapsin and PSD95 respectively (Fig 5D). Probing with anti-KCTD5 antibody revealed that Inc orthologs were present and modestly enriched in synaptosomes with respect to total membrane fractions (Fig 5D). Similarly, endogenous Cul3 was present and enriched in synaptic fractions in its native and higher-molecular weight neddylated form, indicating that active Nedd8-conjugated Cul3 complexes are present at synapses (Fig 5D). Thus, Inc orthologs and Cul3 are present endogenously at mammalian synapses in vivo, suggesting that they form functional ubiquitin ligase complexes at synapses and that their substrates include synaptic proteins.

### Insomniac localizes to the neuronal cytosol, arborizations, and synapses in vivo

To assess the localization of Inc in vivo and determine whether it is similar to that of its mouse orthologs, we examined flies expressing Myc-Inc and 3×FLAG-Inc, forms of Inc which fully rescue the sleep defects of *inc* mutants (Fig 4 and data not shown) and are thus likely to recapitulate attributes of endogenous Inc including its subcellular localization. In the adult brain, expression of 3×FLAG-Inc under the control of *inc-Gal4* yielded strong anti-FLAG signal in cell bodies and in projections including those of the mushroom bodies, ellipsoid body, and fan-shaped body (Fig 6A). To assess the subcellular localization of Inc in adult neurons more clearly, we first identified sparse neuronal populations likely to express *inc* natively, as indicated by their expression of the *inc-Gal4* driver that fully rescues *inc* mutants (Fig 4D). Animals bearing *inc-Gal4* and a nuclear localized GFP reporter (*UAS-nls-GFP*) exhibited GFP signal in corazonin positive (CRZ^+^) neurons in the dorsal brain [34] and in pigment dispersing factor-expressing (PDF^+^) circadian pacemaker neurons [35] (Fig 6B and 6C). We then utilized Gal4 drivers specific for these neuronal subpopulations to express Myc-Inc and assess its localization. In both populations, Myc-Inc was largely excluded from the nucleus and present in neuronal cell bodies and arborizations (Fig 6D and 6F). To assess the nature of projections containing Inc, we compared the pattern of Inc localization to that of the pre-and post-synaptic markers synaptotagmin-GFP (Syt-eGFP) [36] and DenMark [37] expressed in the same neuronal populations. In corazonin neurons, whose presynaptic and dendritic compartments are well separated, Myc-Inc trafficked both to medial dendritic structures and to lateral puncta located in the same regions as presynaptic termini of these neurons (Fig 6D and 6E). In PDF^+^ neurons, Myc-Inc localized to ipsilateral ventral projections to accessory medulla that are of a largely dendritic character (Fig 6F and Fig 6G) [38], but was not detectable in axonal projections to the dorsal brain. Thus, Inc is able to traffic to both pre-and postsynaptic compartments in adult neurons in a manner that may be influenced by cell-type specific factors.

**Fig 6.**
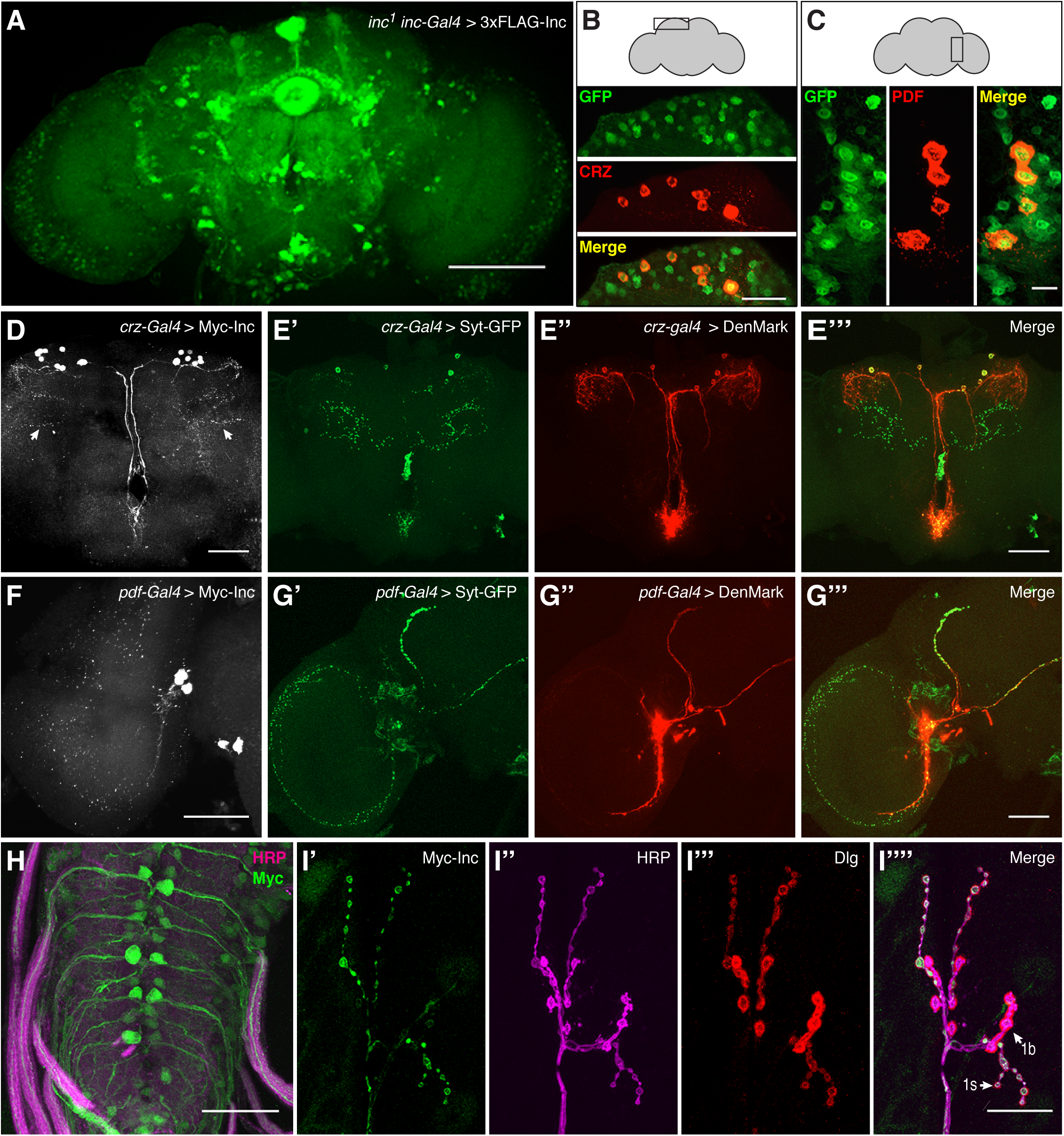
Neuronal localization of Insomniac in vivo. (**A-G**) Immunohistochemical analysis of adult brains. **(A)** Maximal Z-projection of *inc*^*1*^ *inc-Gal4; UAS-3×FLAG-Inc / +* adult brain stained with anti-FLAG. **(B and C)** *inc-Gal4 ; UAS-nls-GFP / +* adult brains stained with anti-GFP, anti-CRZ, and anti-PDF antibodies as indicated. **(D)** *crz-Gal4 / + ; UAS-Myc-Inc / +* brain stained with anti-Myc. Arrows indicate likely presynaptic puncta. **(E)** *crz-Gal4 /+; UAS-DenMark UAS-Syt-eGFP* / + brain stained with anti-GFP and anti-dsRed antibodies. **(F)** *pdf-Gal4; UAS-Myc-Inc / +* brain stained with anti-Myc. **(G)** *pdf-Gal4 ; UAS-DenMark UAS-Syt-eGFP* / + brain stained with anti-GFP and anti-dsRed antibodies. **(H and I)** *inc-Gal4 / + ; UAS-Myc-Inc / +* third instar larval ventral ganglion (H) and NMJ 6/7 (I) stained with anti-Myc, anti-HRP, andanti-Dlg as indicated. Scale bars represent 100 µm in (A), 25 µm in (B), 10 µm in (C), 50 µm in (D-H), and 25 µm in (I).

We also assessed Myc-Inc localization in the third instar larval brain and at the larval neuromuscular junction (NMJ), the latter which permits higher resolution analysis of synaptic termini [39]. In animals bearing *inc-Gal4* and *UAS-Myc-Inc*, we observed Inc signal in motor neuron cell bodies and their axonal projections innervating the NMJ (Fig 6H). As in the adult brain (Fig 6A) [11], Inc expression was present in a subset of neurons, as evident in a fraction of Myc-and HRP-positive projections emanating from the ventral ganglion. At the NMJ itself, Inc was enriched at synaptic boutons and was more prominently expressed within type Is boutons (Fig 6I and S6A Fig). The presence of Inc in motor neurons, their axonal projections, and at boutons circumscribed by postsynaptic DLG signal indicates that Inc signal in these preparations is presynaptic (Fig 6I). Muscle nuclei exhibited weaker Inc signal (Fig 6I), suggesting, along with analysis of additional *inc-Gal4* transgenes (S6B Fig), that *inc* may also be expressed postsynaptically at the NMJ. Taken together with findings that Inc orthologs traffic to mammalian synapses and have functions conserved with those of Inc (Figs 4 and 5), these data suggest that Inc family members may have evolutionarily conserved functions at synapses.

### Altered synaptic anatomy and reduced synaptic strength of *inc* mutants suggests that Inc family proteins may link synaptic function and sleep

A prominent hypothesis invokes synaptic homeostasis as a key function of sleep [40], and findings from vertebrates and flies support the notion that sleep modulates synaptic structure [e.g. 41-44]. Neuronal *inc* activity is essential for normal sleep [11,12], but whether *inc* impacts synaptic function is not known. To test whether the distribution of Inc at synaptic termini reflects a synaptic function, we assessed the anatomical and physiological properties of *inc* mutants at the NMJ. *inc*^*1*^ and *inc*^*2*^ null mutants both exhibited significantly increased bouton number with respect to wild-type animals (Fig 7A and Fig 7B), indicating that Inc is essential for regulation of synaptic growth or plasticity. To assess whether these anatomical defects are associated with altered synaptic transmission, we recorded postsynaptically from muscle in control and transheterozygous *inc*^*1*^*/inc*^*2*^ animals. While the amplitude of spontaneous miniature postsynaptic potentials was not significantly altered in *inc* mutants, their frequency was reduced (Fig 7C-7E). The amplitude of evoked postsynaptic potentials triggered by presynaptic stimulation was significantly reduced in *inc*^*1*^*/ inc*^*2*^ mutants, and quantal content was similarly decreased (Fig 7F-7H). The attenuation of evoked potentials and increased bouton number in *inc* mutants suggest that a compensatory increase in synaptic growth may arise in response to defects in synaptic transmission, though this increase does not compensate for the decreased strength of *inc* synapses. These data indicate that *inc* is vital for normal synaptic structure and physiology and suggest, together with the ability of Inc and its orthologs to localize to synapses, that Inc family members may direct the ubiquitination of proteins critical for synaptic function.

**Fig 7.**
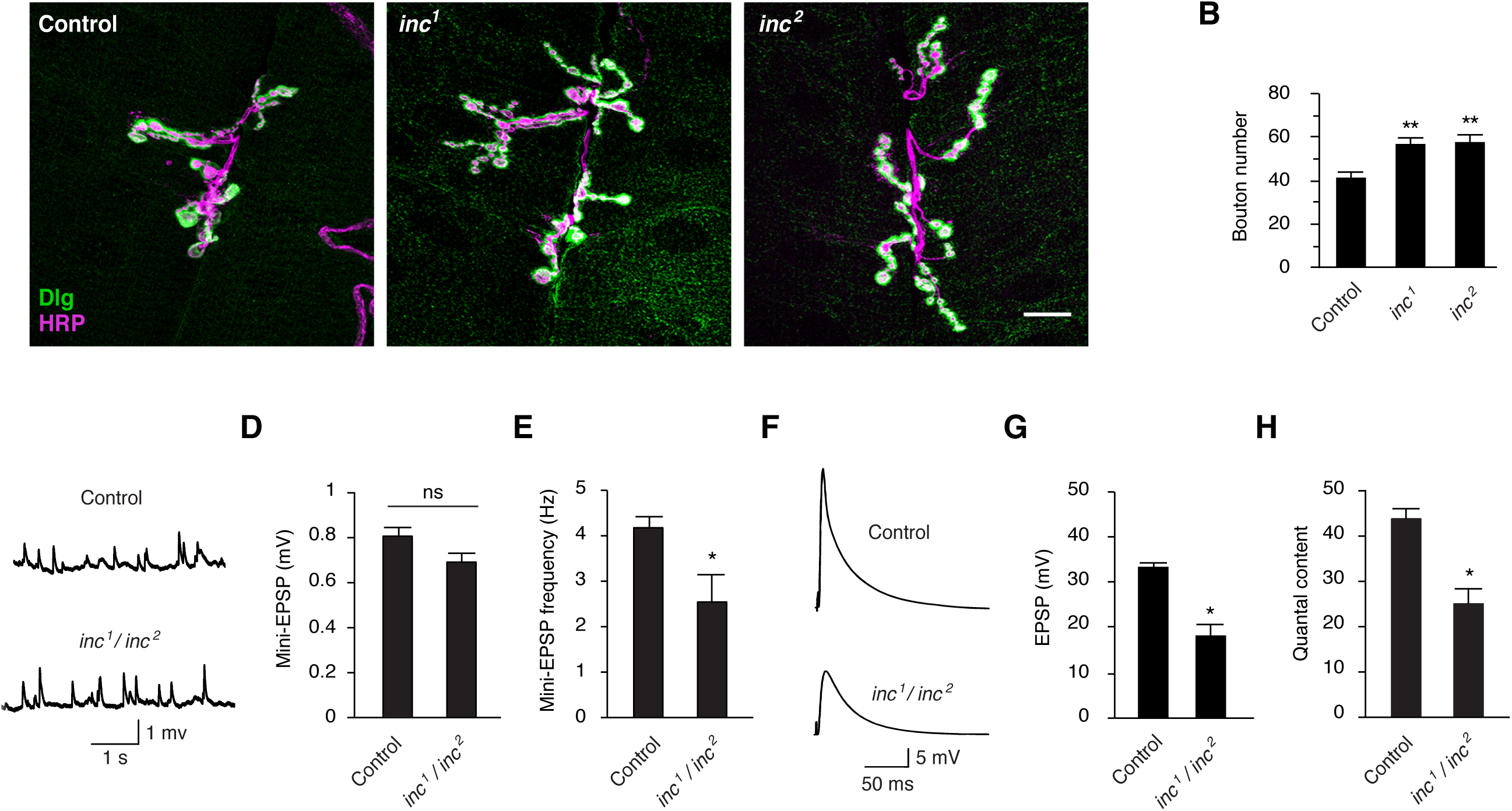
Insomniac is essential for synaptic anatomy and function. **(A)** Female third instar larval abdominal segment 3 muscle 6/7 NMJ are shown from *inc* mutant or isogenic control *w*^*1118*^ animals. Scale bar is 10 µm. **(B)** Total bouton count for indicated genotypes at third instar larval abdominal segment 3 muscle 6/7 NMJ. **(C)** Representative traces of miniature EPSPs for control *w*^*1118*^ and *inc*^*1*^*/inc*^*2*^ animals. **(D)** mEPSP amplitude. **(E)** mEPSP frequency. **(F)** Representative traces of evoked potentials. **(G)** EPSP amplitude. **(H)** Quantal content. For (B), n=40-43, ** p < 0.001 with respect to control. For (D, E, G, and H), n = 11-31, * p < 0.01; ns, not significant (p > 0.05). For all charts, mean ± SEM is shown.

## Discussion

The presence of sleep states in diverse animals has been suggested to reflect a common purpose for sleep and the conservation of underlying regulatory mechanisms [45]. Here we have shown that attributes of the Insomniac protein likely to underlie its impact on sleep in *Drosophila*—its ability to function as a multimeric Cul3 adaptor and engage neuronal targets that impact sleep—are functionally conserved in its mammalian orthologs. Our comparative analysis of Inc family members in vertebrate and invertebrate neurons furthermore reveals that these proteins traffic to synapses and that Inc itself is essential for normal synaptic structure and excitability. These findings support the hypothesis that Inc family proteins serve as Cul3 adaptors and direct the ubiquitination of conserved neuronal substrates that impact sleep and synaptic function.

The ability of KCTD2 and KCTD5 to substitute for Inc in the context of sleep is both surprising and notable given the complexity of sleep-wake behavior and the likely functions of these proteins as Cul3 adaptors. Adaptors are multivalent proteins that self-associate, bind Cul3, and recruit substrates, and these interactions are further regulated by additional post-translational mechanisms [46]. Our findings indicate that KCTD2 and KCTD5 readily substitute for Inc within oligomeric Inc-Cul3 complexes, and strongly suggest that these proteins recapitulate other aspects of Inc function in vivo including the ability to engage neuronal targets that impact sleep. The simplest explanation for why KCTD2 and KCTD5 have retained the apparent ability to engage Inc targets despite the evolutionary divergence of *Drosophila* and mammals is that orthologs of Inc targets are themselves conserved in mammals. This inference draws support from manipulations of *Drosophila* Roadkill/HIB and its mammalian ortholog SPOP, Cul3 adaptors of the MATH-BTB family that regulate the conserved Hedgehog signaling pathway [47]. While the ability of SPOP to substitute for HIB has not been assessed by rescue at an organismal level, clonal analysis in *Drosophila* indicates that ectopically expressed mouse SPOP can degrade the endogenous HIB substrate Cubitus Interruptus (Ci), and conversely, that HIB can degrade mammalian Gli proteins that are the conserved orthologs of Ci and substrates of SPOP [47]. By analogy, Inc targets that impact sleep are likely to have orthologs in vertebrates that are recruited by KCTD2 and KCTD5 to Cul3 complexes. While our manipulations do not resolve whether KCTD17 can substitute for Inc in vivo, the ability of KCTD17 to assemble with fly Inc and Cul3 suggests that any functional divergence among mouse Inc orthologs is likely to arise outside of the BTB domain, and in particular may reflect properties of their C-termini including the ability to recruit substrates.

The finding that Inc transits to synapses and is required for normal synaptic function is intriguing in light of hypotheses that invoke synaptic homeostasis as a key function of sleep [40]. While ubiquitin-dependent mechanisms contribute to synaptic function and plasticity [48-51] and sleep is known to influence synaptic remodeling in both vertebrates and invertebrates [41,43,44,52], molecular links between the two processes remain elusive. Our findings suggest that the neuronal requirement of Inc for normal sleep-wake cycles may be intimately linked to its function at synapses. The synaptic phenotypes of *inc* mutants—increased synaptic growth, decreased evoked neurotransmitter release, and modest effects on spontaneous neurotransmission—are qualitatively distinct from those of other short sleeping mutants. *Shaker* (*Sh*) and *Hyperkinetic* (*Hk*) mutations decrease sleep in adults [26,53] but increase both excitability and synaptic growth at the NMJ [54-56], suggesting that synaptic functions of Inc may affect sleep by a mechanism different than broad neuronal hyperexcitability. While a parsimonious model is that Inc directs the ubiquitination of a target critical for synaptic transmission both at the larval NMJ and in neuronal populations that promote sleep, the precise mechanisms await the elucidation of Inc targets in the context of synaptic physiology and sleep-wake behavior. The identification of Inc targets is also essential to distinguish possible presynaptic and postsynaptic functions of Inc, and whether Inc engages local synaptic proteins or extrasynaptic targets that ultimately influence synaptic function.

A clear implication of our findings is that neuronal targets and synaptic functions of Inc may be conserved in other animals. While the impact of Inc orthologs on sleep in vertebrates is as yet unknown, findings from *C.elegans* support the notion that conserved molecular functions of Inc and Cul3 may underlie similar behavioral outputs in diverse organisms. INSO-1/C52B11.2, the only *C.elegans* ortholog of Inc, interacts with Cul3 [14], and RNAi against Cul3 and INSO-1 reduces the duration of lethargus, a quiescent sleep-like state, suggesting that effects of Cul3-and Inc-dependent ubiquitination on sleep may be evolutionarily conserved [57]. The functions of Inc orthologs and Cul3 in the mammalian nervous system await additional characterization, but emerging data suggest functions relevant to neuronal physiology and disease. Human mutations at the *KCTD2/ACGH* locus are associated with Alzheimer’s disease [58], and mutations of *KCTD17* with myoclonic dystonia [59]. *Cul3* lesions have been associated in several studies with autism spectrum disorders [60-62] and comorbid sleep disturbances [62]. More generally, autism spectrum disorders are commonly associated with sleep deficits [63] and are thought to arise in many cases from altered synaptic function [64], but molecular links to sleep remain fragmentary. Studies of Inc family members and their conserved functions in neurons are likely to broaden our understanding of how ubiquitination pathways may link synaptic function to the regulation of sleep and other behaviors.

## Methods

### RT-PCR and in situ hybridization

Total RNA was isolated with TRIZOL (ThermoFisher) from a single brain hemisphere of a mixed C57BL/6 background mouse. 5 µg total RNA was annealed to random hexamer primers and reverse transcribed with Thermoscript (ThermoFisher) according to the manufacturer’s protocol. KCTD2, KCTD5, and KCTD17 transcripts were amplified using primer pairs oNS286 and oNS287, oNS288 and oNS289, and oNS290 and oNS291, respectively.

For in situ hybridization, DNA templates bearing a terminal SP6 promoter for in vitro transcription were generated by PCR amplification of C57BL/6 mouse genomic DNA, using primer pairs oNS1204 and oNS1205 for KCTD2, oNS1207 and oNS1208 for KCTD5, and oNS1213 and oNS1214 for KCTD17. Riboprobes were transcribed with SP6 polymerase and DIG-11-UTP or Fluorescein-12-UTP (Roche). In situ hybridization was performed as described [65], amplifying Fluorescein-and DIG-labeled probes with Fluorescein-tyramide and Cy5-tyramide (Perkin Elmer) respectively.

### Plasmids and molecular cloning

Vectors for Drosophila transgenesis were as follows: pUASTattB-Myc-Inc (pNS346) encodes a 1× N-terminal Myc epitope (MEQKLISEEDLAS) fused to Inc, and was generated by three piece ligation of BglII-XhoI digested pUASTattB [66], a BglII-NheI 1×Myc fragment generated by phosphorylating and annealing oligonucleotides oNS283 and oNS284, and an NheI-XhoI *inc* fragment liberated from the PCR amplification product of pUAST-Inc (pNS272) [11] template and primers oNS277 and oNS285.

pUASTattB-Myc-KCTD2 (pNS347) was generated similarly to pNS346, substituting a NheI-XhoI KCTD2 fragment liberated from the PCR amplification product of mouse brain cDNA and primers oNS286 and oNS287. Amplified KCTD2 sequences are identical to those within GenBank accession NM_183285.3.

pUASTattB-Myc-KCTD5 (pNS348) was generated similarly to pNS346, substituting a NheI-XhoI KCTD5 fragment liberated from the PCR amplification product of mouse brain cDNA and primers oNS288 and oNS289. Amplified KCTD5 sequences are identical to those within GenBank accession NM_027008.2.

pUASTattB-Myc-KCTD17.3 (pNS349) and pUASTattB-Myc-KCTD17.2 (pNS350) were generated similarly to pNS346, substituting NheI-XhoI KCTD17 fragments liberated from the PCR amplification products of mouse brain cDNA and primers oNS290 and oNS291. The smaller and larger NheI-XhoI fragments respectively corresponding to KCTD17.3 and KCTD17.2 were gel purified and ligated separately. Amplified KCTD17.3 and KCTD17.2 sequences are identical to those within GenBank accession NM_001289673.1 and NM_001289672.1 respectively.

pUAST-Inc-HA (pNS273) encodes Inc fused to a 1× C-terminal HA epitope (GSYPYDVPDYA) and was generated by three piece ligation of pUAST BglII-XhoI, a BglII-EcoRI *inc* fragment liberated from pNS272 [11], and an EcoRI-XhoI HA fragment generated by phosphorylating and annealing oligonucleotides oNS191 and oNS192.

pUASTattB-3×FLAG-Inc (pNS404) encodes a 3× N-terminal FLAG epitope (MDYKDDDDKGSDYKDDDDKGSDYKDDDDKAS) fused to Inc and was generated by three piece ligation of EcoRI-XhoI digested pUASTattB, an EcoRI-NheI 3×FLAG fragment liberated from the PCR amplification product of pNS311 template and primers ACF and oNS241, and a NheI-XhoI *inc* fragment liberated from pNS351.

Vectors for expression in S2 cells were as follows:

pAc5.1-3×Myc-Inc (pNS351) encodes a 3× N-terminal Myc epitope (MEQKLISEEDLGSEQKLISEEDLGSEQKLISEEDLAS) fused to Inc in a derivative of pAc5.1/V5-HisA (ThermoFisher), and was generated by ligating NheI-XhoI digested pNS309 [11] to a NheI-XhoI *inc* fragment prepared as for pNS346.

pAc5.1-3×Myc-KCTD2 (pNS352) was generated similarly to pNS351, substituting a NheI-XhoI KCTD2 fragment prepared as for pNS347.

pAc5.1-3×Myc-KCTD5 (pNS353) was generated similarly to pNS351, substituting a NheI-XhoI KCTD5 fragment prepared as for pNS348.

pAc5.1-3×Myc-KCTD17.3 (pNS391) was generated by ligating NheI-XhoI digested pNS309 to a NheI-XhoI KCTD17.3 fragment liberated from the PCR amplification product of pNS354 template and primers oNS290 and oNS612. pNS354 was generated by ligating NheI-XhoI digested pNS309 and the NheI-XhoI KCTD17.3 fragment prepared as for pNS349.

pAc5.1-3×Myc-KCTD17.2 (pNS392) was generated by ligating NheI-XhoI digested pNS309 to a NheI-XhoI KCTD17.2 fragment liberated from the PCR amplification product of pNS355 template and primers oNS290 and oNS955. pNS355 was generated by ligating NheI-XhoI digested pNS309 and the NheI-XhoI KCTD17.2 fragment prepared as for pNS350.

pAc5.1-3×HA-Inc (pNS402) was generated by ligating NheI-XhoI digested pNS310 [11] and a NheI-XhoI *inc* fragment liberated from pNS351.

pAc5.1-3×HA-mCul3 (pNS367) encodes a 3× N-terminal HA epitope (MYPYDVPDYAGSYPYDVPDYAGSYPYDVPDYAAS) fused to mouse Cul3, and was generated by three piece ligation of NheI-NotI digested pNS310 [11], a 0.3 kb NheI-HindIII 5’ mCul3 fragment liberated from the PCR amplification product of pCMV-SPORT6-mCul3 template (ThermoFisher, GenBank accession BC027304) and primers oNS313 and oNS314, and a 2.2 kb HindIII-NotI 3’ mCul3 fragment generated by digesting pCMV-SPORT6-mCul3 with AvaI, blunting with T4 DNA polymerase, ligating a NotI linker, and digesting with HindIII-NotI.

pAc5.1-3×FLAG-Cul3 (pNS403) encodes a 3× N-terminal FLAG epitope fused to *Drosophila* Cul3 and was generated by ligating NheI-NotI digested pNS311 and a NheI-NotI Cul3 fragment liberated from pNS314 [11]. pNS311 contains a N-terminal 3×FLAG tag and was generated from pNS298, a derivative of pAc5.1/V5-HisA that contains a C-terminal 3×FLAG tag. To construct pNS298, oligonucleotides oNS234 and oNS235 were phosphorylated, annealed, and cloned into XhoI-XbaI digested pAc5.1/V5-HisA. To construct pNS311, the EcoRI-NotI fragment liberated from the PCR amplification product of pNS298 template and primers oNS240 and oNS241 was ligated to EcoRI-NotI digested pAc5.1/V5-HisA.

pAc5.1-Inc-GFP (pNS275) encodes Inc fused at its C-terminus to GFP and was generated by three piece ligation of EcoRI-NotI digested pAc5.1/V5-HisA, an EcoRI-BamHI *inc* fragment liberated from pNS273, and a BamHI-NotI EGFP fragment liberated from pEGFP-N3 (Clontech).

Mammalian expression vectors were as follows:

pcDNA3.1-3×Myc-Inc (pNS358) was generated by ligating NheI-XhoI digested pcDNA3.1(+) (ThermoFisher) with a SpeI-XhoI 3×Myc-Inc fragment liberated from pNS351.

pcDNA3.1-3×Myc-KCTD2 (pNS359) was generated similarly to pNS358, substituting a SpeI-XhoI 3×Myc-KCTD2 fragment liberated from pNS352.

pcDNA3.1-3×Myc-KCTD5 (pNS360) was generated by ligating EcoRI-XhoI digested pcDNA3.1(+) with a EcoRI-XhoI 3×Myc-KCTD5 fragment liberated from pNS353.

pcDNA3.1-3×Myc-KCTD17.3 (pNS393) was generated by three piece ligation of EcoRI-XhoI digested pcDNA3.1(+), an EcoRI-NheI 3×Myc fragment liberated from the PCR amplification product of pNS351 template and primers ACF and oNS318, and a NheI-XhoI KCTD17.3 fragment liberated from the PCR amplification product of pNS354 template and primers oNS290 and oNS612.

pcDNA3.1-3×Myc-KCTD17.2 (pNS394) was generated similarly to pNS393, substituting a NheI-XhoI KCTD17.2 fragment liberated from the PCR amplification product of pNS355 template and primers oNS290 and oNS955.

pcDNA3.1-3×HA-Cul3 (pNS365) was generated by ligating EcoRI-NotI digested pcDNA3.1(+) with the EcoRI-NotI 3×HA-Cul3 fragment from pNS314 [11].

pcDNA3.1-3×HA-mCul3 (pNS369) was generated by ligating KpnI-NotI digested pcDNA3.1(+) with the KpnI-NotI 3×HA-mCul3 fragment liberated from pNS367.

pcDNA3.1-3×FLAG-Inc (pNS395) encodes a 3× N-terminal FLAG epitope fused to Inc, and was generated by three piece ligation of EcoRI-XhoI digested pcDNA3.1(+), an EcoRI-NheI 3×FLAG fragment liberated from the PCR amplification product of pNS386 template and primers ACF and oNS318, and the NheI-XhoI *inc* fragment liberated from pNS351. pNS386 was generated by ligating NheI-XhoI digested pNS311 and an NheI-XhoI *inc* fragment liberated from pNS351.

pcDNA3.1-3×FLAG-KCTD2 (pNS396) was generated similarly to pNS395, substituting an NheI-XhoI KCTD2 fragment liberated from pNS352.

pcDNA3.1-3×FLAG-KCTD5 (pNS397) was generated similarly to pNS395, substituting an NheI-XhoI KCTD5 fragment liberated from pNS353.

pcDNA3.1-3×FLAG-KCTD17.2 (pNS398) was generated similarly to pNS395, substituting the NheI-XhoI fragment liberated from the PCR amplification product of pNS355 template and primers oNS290 and oNS955.

pcDNA3.1-3×FLAG-KCTD17.3 (pNS399) was generated similarly to pNS395, substituting the NheI-XhoI fragment liberated from the PCR amplification product of pNS354 template and primers oNS290 and oNS612.

pcDNA3-Inc (pNS279) encodes Inc fused at its C-terminus to GFP and was generated by ligating EcoRI-BamHI digested pEGFP-N3 (Clontech) and an EcoRI-BamHI *inc* fragment liberated from pNS273.

### Oligonucleotides

Oligonucleotides used in this work, listed 5’ to 3’, are as follows:

oNS119: GTCCGCGCGATTCCCTTGCTTGC
oNS191: AATTTTGGGAATTGGATCCTACCCCTACGATGTGCCCGATTACGCCTAAC
oNS192: TCGAGTTAGGCGTAATCGGGCACATCGTAGGGGTAGGATCCAATTCCCAA
oNS198: ACTGGGATCCATCCGCCTGTGTGGCTGGGACGG
oNS234: TCGAGGCTAGCGACTACAAGGATGATGACGATAAGGGCTCCGATTACAAGGACGACGATGATAAGGGATCCGATTACAAGGATGATGACGACAAGTGAT
oNS235: CTAGATCACTTGTCGTCATCATCCTTGTAATCGGATCCCTTATCATCGTCGTCCTTGTAATCGGAGCCCTTATCGTCATCATCCTTGTAGTCGCTAGCC
oNS240: ACTGGAATTCCGCGGCAACATGGACTACAAGGATGATGACGATAAGGGC
oNS241: ACTGGCGGCCGCTCCTAGGGTGCTAGCCTTGTCGTCATCATCCTTGTAATCGGAT
oNS254: GTGCGCCAAGTGTCTGAAGAACAACTGG
oNS255: GATGAGCTGCCGAGTCAATCGATACAGTC
oNS277: ACGTGCTAGCATGAGCACGGTGTTCATAAACTCGC
oNS283: GATCTCAACATGGAGCAGAAGCTGATCAGCGAGGAGGATCTGG
oNS284: CTAGCCAGATCCTCCTCGCTGATCAGCTTCTGCTCCATGTTGA
oNS285: ACGTGCTAGCTCGAGGGGTTGTGTGTGAATATATAGCGCGA
oNS286: ACGTGCTAGCATGGCGGAGCTGCAGCTGG
oNS287: ACGTGCTAGCTCGAGCCGCTTACATTCGAGAGCCTCTCTCC
oNS288: ACGTGCTAGCATGGCGGAGAATCACTGCGAGCTG
oNS289: ACGTGCTAGCTCGAGCCTCACATCCTTGAGCCCCGTTC
oNS290: ACGTGCTAGCATGCAGACAACGCGGCCGGCG
oNS291: ACGTGCTAGCTCGAGCCCAAGGCAGGAGTGAGTCTCAGC
oNS313: ACGTGCTAGCATGTCGAATCTGAGCAAAGGCACGGG
oNS314: GCCGAAGATGATCCCTAATACACCCATACCG
oNS318: ACGTCTCGAGTTACTGCGTCACGTTGTAGAACTC
oNS612: ACGTCTCGAGTCACATCCGGGTGCCTCTGGCTT
oNS955: ACGTCTCGAGTCACTGCAAGCTCAGGCTTGGGTCTG
oNS1204: TAATACGACTCACTATAGGGGAAGGCAAGAGAGCAATCGGC
oNS1205: GCGATTTAGGTGACACTATAGAAGAAAAGGCTGCAGAAGCAGTTAC
oNS1207: TAATACGACTCACTATAGGGGGCTCAAGGATGTGAGGAATGCTGAG
oNS1208: GCGATTTAGGTGACACTATAGAAGCAGCCTCTATCCCAGGCACAAC
oNS1213: TAATACGACTCACTATAGGGTTACAAGCCAGAGGCACCCGGA
oNS1214: GCGATTTAGGTGACACTATAGAAGCAGCTCAACCCGTTACACCTGTC
ACF: GACACAAAGCCGCTCCATCAG
attP2-5’: CACTGGCACTAGAACAAAAGCTTTGGCG

### Cell culture and biochemistry

293T cells were cultured in DMEM containing 10% FBS, penicillin, and streptomycin, and transfected with Lipofectamine 2000 (ThermoFisher) according to the manufacturers protocol. S2 cells were cultured in S2 media containing 10% FBS, penicillin, and streptomycin, and were transfected with Effectene (Qiagen) as described previously [11]. For both 293T and S2 cells, transfections were performed in 6 well or 12 well plates for ~24h until liposome-containing media was replaced with fresh culture media. Cells or coverslips were harvested for lysis or immunohistochemistry 36-48h after transfections were initiated. For transfections involving more than one plasmid, an equal amount of each was used. Rat cortical neurons were cultured on poly-D-lysine coated coverslips and transfected with calcium phosphate at 7 days in vitro (DIV) as described previously [67].

For co-immunoprecipitations from 293T cells, samples were lysed with ice-cold RIPA buffer (50 mM Tris-Cl pH 7.6, 150 mM NaCl, 50 mM NaF, 2 mM EDTA, 0.5% sodium deoxycholate, 1% NP40, 0.1% SDS) containing protease inhibitor (Sigma, P8340). For co-immunoprecipitation of Inc family members from S2 cells, samples were lysed with ice-cold NP40 buffer (50 mM Tris pH 7.6, 150mM NaCl, 0.5% NP40) containing protease inhibitors or RIPA buffer as above; for S2 cell co-immunoprecipitations involving Cul3, ice-cold NP40 buffer was used. Protein extracts were quantitated in duplicate (BioRad, 5000111) and 160-400 µg (293T) or 800-1000 µg (S2) was immunoprecipitated with 20 µl (50% slurry) of anti-FLAG (Sigma, F2426) or anti-HA (Sigma, E6779) affinity gel for 1 hr nutating at 4**°**C. Samples were then washed 4×5 min at 4**°**C with lysis buffer, denatured in SDS sample buffer, separated on Tris SDS-PAGE gels, and transferred to nitrocellulose. Membranes were blocked for 1 hr at room temperature or 4**°**C overnight in LI-COR Odyssey buffer (LI-COR, 927-40000) or 1% casein in PBS. Membranes were subsequently incubated in blocking buffer containing 0.1% Tween 20 and the appropriate primary antibodies: rabbit anti-Myc (1:2,000, Sigma, C3956), mouse anti-FLAG (1:2,000, Sigma, F1804), rat anti-HA (1:1,000-1:2,000, Roche, 11867431001), rabbit anti-Cul3 (1:1,000, Bethyl Laboratories, A301-109A), and rabbit anti-KCTD5 (1:2,000, Proteintech, 15553-1-AP). After washing 4×5 min in a solution containing 150 mM NaCl, 10mM Tris pH 7.6, and 0.1% Tween 20 (TBST), membranes were incubated in the dark for 30 min at room temperature with appropriate secondary antibodies, all diluted 1:30,000 in blocking buffer containing 0.1% Tween 20 and 0.01% SDS: Alexa 680 donkey anti-rabbit (Life Technologies, A10043), Alexa 790 anti-mouse (Life Technologies, A11371), and Alexa 790 anti-rat (Jackson ImmunoResearch, 712-655-153). Membranes were then washed 4×5 min in TBST, 1×5 min in TBS, and imaged on a Li-Cor Odyssey CLx instrument.

Fly protein extracts were prepared from whole animals or from sieved heads by manual pestle homogenization in ice-cold NP40 lysis buffer supplemented with protease inhibitors. 50 µg was separated on Tris-SDS-PAGE gels and blotted as described above. Primary antibodies were mouse anti-Myc (1:1,000, BioXCell, BE0238; or 1:1,000, Cell Signaling Technology, 2276), rabbit anti-tubulin (1:30,000, VWR, 89364-004), or mouse anti-actin (1:1,000, Developmental Studies Hybridoma Bank (DSHB), JLA20). Secondary antibodies were Alexa 680 donkey anti-rabbit, Alexa 680 anti-mouse (Life Technologies, A10038), and Alexa 790 anti-rabbit (Life Technologies, A11374).

Mouse and rat brain extracts not used for synaptosome preparations were prepared by homogenizing brains in ice-cold NP40 lysis buffer supplemented with protease inhibitors. Extracts were separated with SDS-PAGE and blotted as described above.

Synaptosomes were prepared from rat brain essentially as described [68], and probed with rabbit anti-Cul3 (1:1,000), mouse anti-Actin (1:1,000), rabbit anti-KCTD5 (1:2,000), guinea-pig anti-synapsin (1:1,000, Synaptic Systems, 106 004), and mouse anti-PSD95 (1:1,000, Neuromab, 75-028). Alexa 680 and Alexa 790 secondary antibodies were used as described above.

### Immunohistochemistry and microscopy

Rat cortical neurons transfected at 7 DIV were processed at 13 DIV for immunohistochemistry. Samples were fixed for 15 min with ice-cold 4% paraformaldehyde in Dulbecco’s PBS containing 4 mM EGTA and 4% sucrose, and subsequently permeabilized for 10 min in PBS containing 0.5% normal donkey serum (Lampire Biological, 7332100) and 0.1% Triton X-100. Samples were then blocked at room temperature for 30 min in PBS containing 7.5% normal donkey serum and 0.05% Triton X-100, incubated overnight at 4**°**C in primary antibody cocktail prepared in PBS containing 5% normal donkey serum and 0.05% Triton X-100, and washed 3×5 min in PBS at room temperature. Secondary antibody cocktails were prepared similarly and incubated with samples at room temperature for 30-40 min in the dark. Samples were then washed 3×5 min in PBS at room temperature and mounted on microscope slides in Vectashield (Vector Labs, H-1000). Primary antibodies were rabbit anti-Myc (1:200), mouse anti-PSD95 (1:1,000), and guinea pig anti-synapsin (1:1,000). Secondary antibodies, all diluted at 1:1,000, were Alexa 488 donkey anti-rabbit (Life Technologies, A21206), Alexa 488 donkey anti-guinea pig (Jackson ImmunoResearch, 706-545-148), Alexa 568 donkey anti-mouse (Life Technologies, A10037), and Alexa 568 donkey anti-rabbit (Life Technologies, A10042).

Immunohistochemistry for 293T cells was performed similarly; cells were plated on poly-L-lysine treated coverslips and cultured and transfected as described above. 0.4 µg/ml DAPI was included in the penultimate wash prior to mounting coverslips on microscope slides.

For immunohistochemistry of adult fly brains, whole animals were fixed with 4% paraformaldehyde in PBS containing 0.2% Triton X-100 (PBST) for 3 hr at 4**°**C, and subsequently washed 3×15 min at room temperature with PBST. Brains were dissected in ice-cold PBST, incubated for 30 min at room temperature in blocking solution containing PBST and 5% normal donkey serum, and incubated overnight at 4**°**C in primary antibody cocktail diluted in blocking solution. After 3×15 min washes in PBST at room temperature, samples were incubated in secondary antibody cocktail in blocking solution for 1-3 days at 4**°**C, washed 3×15 min at room temperature with PBST, and mounted on microscope slides in Vectashield. Primary antibodies were mouse anti-FLAG (1:100), rabbit anti-GFP (1:3,000; Life Technologies, A11122), mouse anti-GFP (1:1,000, DSHB, G1), mouse anti-PDF (1:500, DSHB, C7), rabbit anti-CRZ (1:1,000, [69]), rabbit anti-DsRed (1:1,000, Clontech, 632496), and mouse anti-Myc (1:100, BioXCell, BE0238). Secondary antibodies, all used at 1:1,000, were Alexa 488 donkey anti-mouse (Life Technologies, A21202), Alexa 488 donkey anti-rabbit, Alexa 568 donkey anti-mouse, and Alexa 568 donkey anti-rabbit.

For immunohistochemistry of larval brains and neuromuscular junctions, wandering third instar larvae were dissected in PBS and pinned to 35mm Sylgard-coated petri dishes. Where experiments required larvae of a specific sex, gonads were identified prior to dissection by visual inspection of animals under PBS immersion as described [70]. Larval filets were fixed for 30 min at room temperature in 4% paraformaldehyde in PBS, and subsequently rinsed twice and washed 3×20 min in PBST. Samples were blocked in 5% normal donkey serum in PBST at room temperature for 30 min and incubated overnight at 4**°**C in primary antibody cocktail diluted in blocking solution. Samples were then washed 3×20 min in PBST at room temperature and incubated overnight at 4**°**C with secondary antibody cocktail in blocking solution, washed, and mounted in Vectashield. Primary antibodies were rabbit anti-myc (1:500), mouse anti-Dlg (1:1,000, DSHB, 4F3), and Alexa 647 goat anti-HRP (1:200, Jackson ImmunoResearch, 123-605-021). Secondary antibodies, used at 1:1000, were Alexa 488 donkey anti-rabbit and Alexa 568 donkey anti-mouse. Neuromuscular junctions were imaged with a confocal microscope and z-stacks were captured using 40× or 63× oil objectives at 512×512 resolution. Boutons were counted offline using a manual tally counter while manipulating z-stacks in 3-dimensional space using Zen software (Zeiss); each axon branch was counted separately to avoid undercounting or duplicate counts and counts were performed three times to ensure consistency. All bouton counting was performed in a double-blind manner with codes revealed after the entire experiment was scored.

For live imaging of Inc-GFP in S2 cells, cells were cultured on poly-L-lysine coated coverslips and transfected with Effectene as described above. 48 hours post-transfection, coverslips were inverted onto a drop of PBS on microscope slides and imaged immediately (within 5 minutes) on a confocal microscope. For imaging of fixed Inc-GFP signal in S2 cells, coverslips were washed twice with PBS, fixed for 25 min in PBS containing 4% paraformaldehyde, washed twice in PBS, and inverted onto drops of Vectashield containing DAPI (Vector) on microscope slides.

All imaging was performed on Zeiss LSM510 or LSM800 confocal microscopes.

### Fly stocks and transgenes

*elav*^*c155*^*-Gal4* [71] and *inc*^*1*^, *inc*^*2*^, and *inc-Gal4* [11] have been described previously. Unless noted otherwise, all experiments were performed with the X-linked *inc-Gal4* transgene. The *inc*^*1*^ *inc-Gal4* stock was generated by meiotic recombination between isogenic *inc*^*1*^ and *inc-Gal4* chromosomes and verified with duplex PCR using primers oNS254 and oNS255 for Gal4 and oNS119 and oNS198 for *inc*^*1*^. pUASTattB-based vectors generated in this study were integrated at *attP2* [25] with phiC31 recombinase (BestGene); integration was verified by PCR using primer attP2-5’ paired with oNS277, oNS286, oNS288, or oNS290 for Inc, KCTD2, KCTD5, and KCTD17 respectively. All transgenes were backcrossed eight generations to Bloomington stock 5905, an isogenic *w*^*1118*^ stock described elsewhere as iso31 [72].

### Sleep analysis

Crosses were set with five virgin females and three males on cornmeal, agar, and molasses food. One to four day old male flies eclosing from LD-entrained cultures raised at 25**°**C were loaded in glass tubes containing cornmeal, agar, and molasses food. Animals were monitored for 5-7 days at 25**°**C in LD cycles using DAM2 monitors (Trikinetics). The first 36-48 hours of data was discarded to permit animals to acclimate to glass tubes, and an integral number of days of data (3-5) were analyzed using custom Matlab software as described previously [11]. Locomotor data were collected in 1 min bins, and sleep was defined by inactivity for 5 minutes or more; a given minute was assigned as sleep if an animal was inactive for that minute and the preceding four minutes. Dead animals were excluded from analysis by a combination of automated filtering and visual inspection of locomotor traces.

### Electrophysiology

Electrophysiological recordings were performed from abdominal segment 3 muscle 6/7 of third instar larvae as described in [73].

### Statistics

One-way ANOVA and Tukey post-hoc tests were used for comparisons of total sleep, daytime sleep, nighttime sleep, sleep bout number, and bouton number. Nonparametric Kruskal-Wallis tests and Dunn’s post hoc tests were used for comparisons of sleep bout length. Unpaired two-sided Student’s t-tests were used for comparisons of all electrophysiological parameters.

### Sequence alignments

Alignments were performed with Clustal Omega 1.2.1 and BOXSHADE. GenBank accession numbers for transcript variants referred to in Figures S1 and S2 are: KCTD2, NM_183285; KCTD5, NM_027008; KCTD17.1, NM_001289671; KCTD17.2, NM_001289672; KCTD17.3, NM_001289673; KCTD17.4, NR_110357; KCTD17.v1, XM_006521460; KCTD17.v2, XM_011245739; KCTD17.v3, XM_011245740; KCTD17.v4, XM_011245741; KCTD17.v5, XM_006521461; KCTD17.v6, XM_006521462; KCTD17.v7, XM_011245742; KCTD17.v8, XM_006521465; hKCTD17.2, NM_024681.

## Author contributions

Q.L. generated constructs and stocks, and performed behavioral assays, transfections, immunoprecipitations, and immunohistochemical analysis; D.A.K. performed transfections, immunoprecipitations, immunohistochemical analysis, and prepared synaptosomes; H.A.M.H. performed behavioral assays and immunohistochemical analysis; T.Y. performed immunohistochemical analysis; S.S. derived cortical cultures and prepared synaptosomes; R.P.M. performed in situ hybridizations; C.A.F. performed all electrophysiological recordings; N.S. generated constructs and stocks, and performed behavioral assays, transfections, immunoprecipitations, and immunohistochemical analysis. All authors contributed to the design and analysis of experiments. N.S wrote the manuscript with contributions from Q.L.

## Acknowledgements

We thank G. Fishell and R. Tsien for sharing facilities; B. Schulman for discussions of Cul3 adaptors; and J. Dasen, N. Ringstad, M. Shirasu-Hiza, J. Treisman, R. Tsien, and members of the Stavropoulos lab for comments on the manuscript and discussions. N.S. thanks M. Young for support during the initiation of this work. This research was supported by an International Student Research Fellowship from the Howard Hughes Medical Institute (Q.L.), by NIH NS062738 and a Whitehall foundation grant (C.A.F.), and by grants from the Whitehall, Alfred P. Sloan, Leon Levy, and Mathers Foundations, a NARSAD Young Investigator Award from the Brain and Behavior Research Foundation, a New York University Whitehead Fellowship, and the J. Christian Gillin, M.D. Research Award from the Sleep Research Society Foundation (N.S.).

## Supporting information

**S1 Fig.**
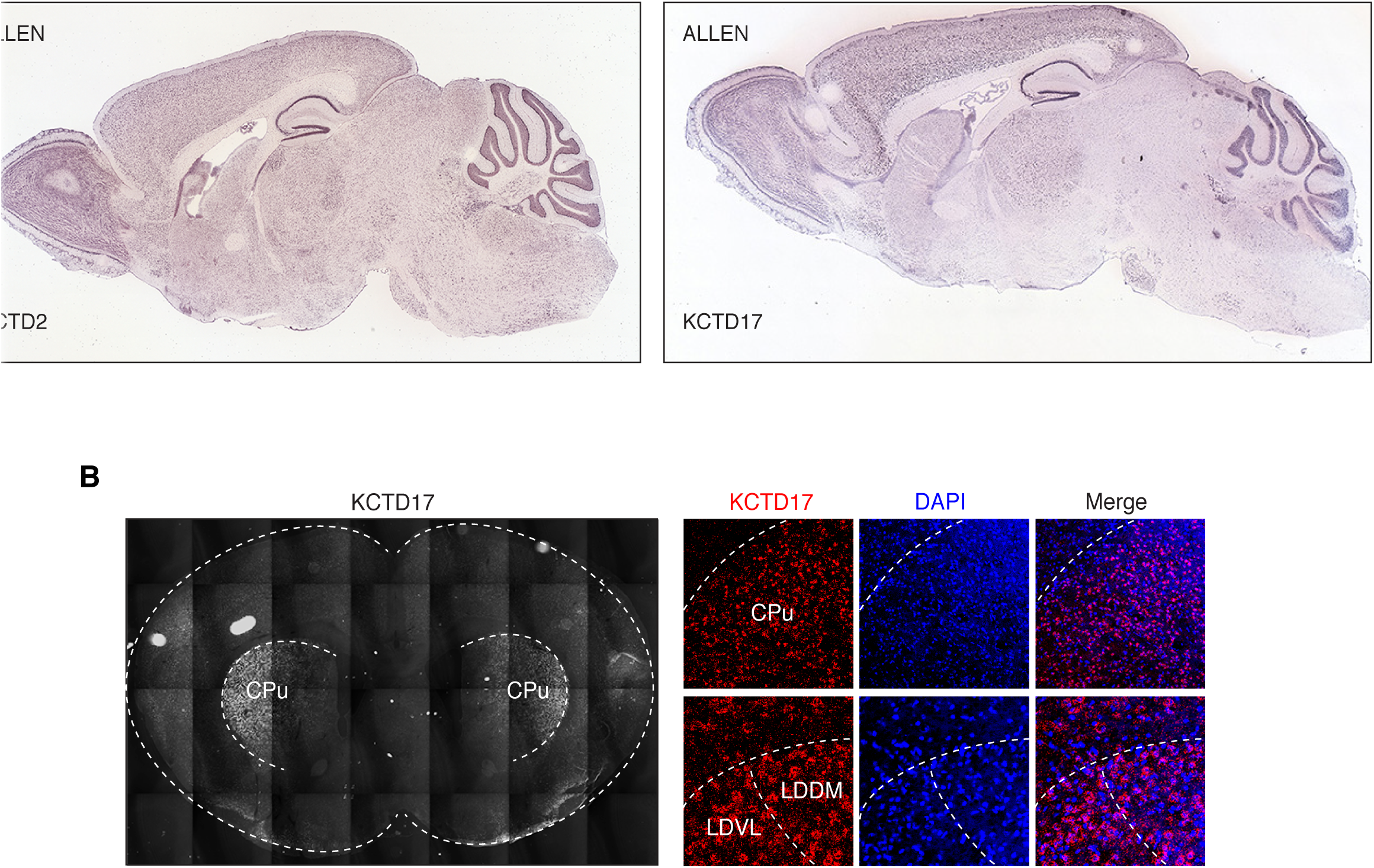
In situ hybridization for mouse Insomniac orthologs. **(A)** Allen Brain Atlas in situ hybridization images for KCTD2 and KCTD17. Brain regions with KCTD2 signal include cortex, hippocampus, striatum, thalamus, hypothalamus, cerebellum, pons, and medulla. KCTD17 signal is more sparse and is present in cortex, hippocampus, striatum, thalamus, and cerebellum. KCTD17 probe in these experiments is complementary to KCTD17 transcript isoforms v1, v2, and v5 (see Fig.S2 and Materials and Methods). **(B)** In situ hybridization of mouse coronal brain section using a probe complementary to KCTD17 transcript isoforms 1, 2, 3, 4, v2, and v4. Signal is prominent in the caudate putamen (CPu) and thalamus, as shown for the laterodorsal thalamic nucleus (LDDM) and laterodorsal ventrolateral thalamus (LDVL), but weak or absent from cortex (left panel and data not shown), suggesting that cortical KCTD17 signal in (A) may reflect differential expression of a nonoverlapping subset of KCTD17 transcript isoforms.

**S2 Fig.**
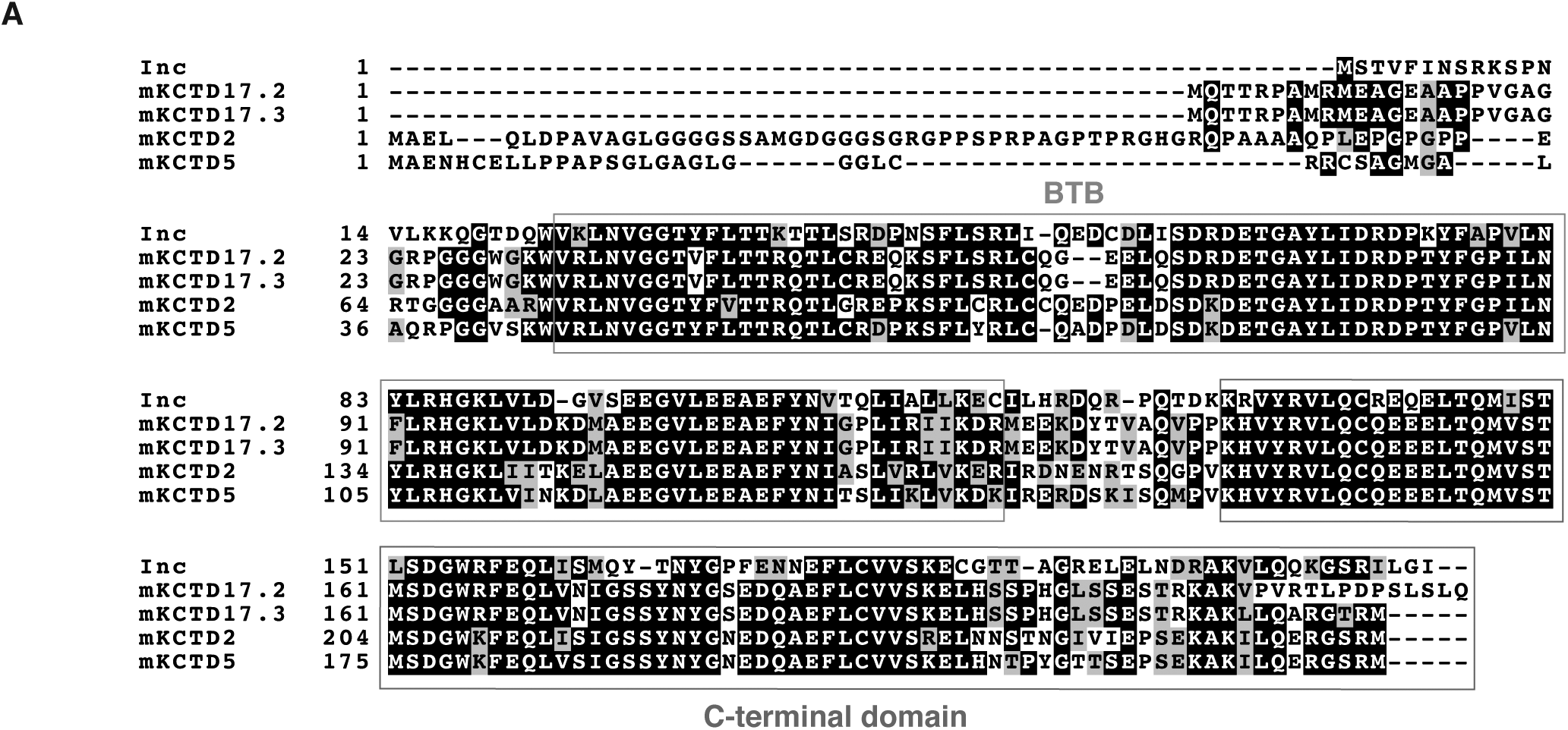
Sequence analysis of mouse Insomniac orthologs. **(A)** Alignment of Inc and its mouse orthologs. Conserved BTB and C-terminal domains are indicated. **(B)** Analysis of alternatively spliced forms of mouse KCTD17 (mKCTD17). Proteins represented by multiple mouse brain cDNA clones in this study are shown alongside additional predicted splice variants from GenBank (see Materials and Methods). The sequence of a KCTD17 protein encoded by human KCTD17 transcript isoform 2 (hKCTD17.2) as characterized in Kasahara et al [24] is shown. Deletion of underlined hKCTD17.2 residues abolishes interaction with trichoplein. Note that residues implicated in binding trichoplein are absent in KCTD2, KCTD5, KCTD17.2, KCTD17.3, and Inc. Alternative splicing of KCTD17 in cloned transcripts and predicted variants occurs after a common exon whose terminal residues are KAK, with color indicating residues shared in different protein isoforms. Proteins encoded by a subset of predicted mKCTD17 transcript variants (v1, v4, v8) are shown; analysis of sequence databases suggests that these alternatively spliced variants may be conserved in humans and other vertebrates. Other predicted mKCTD17 variants not shown (v2, v3, v5, v6, v7) may also be conserved.

**S3 Fig.**
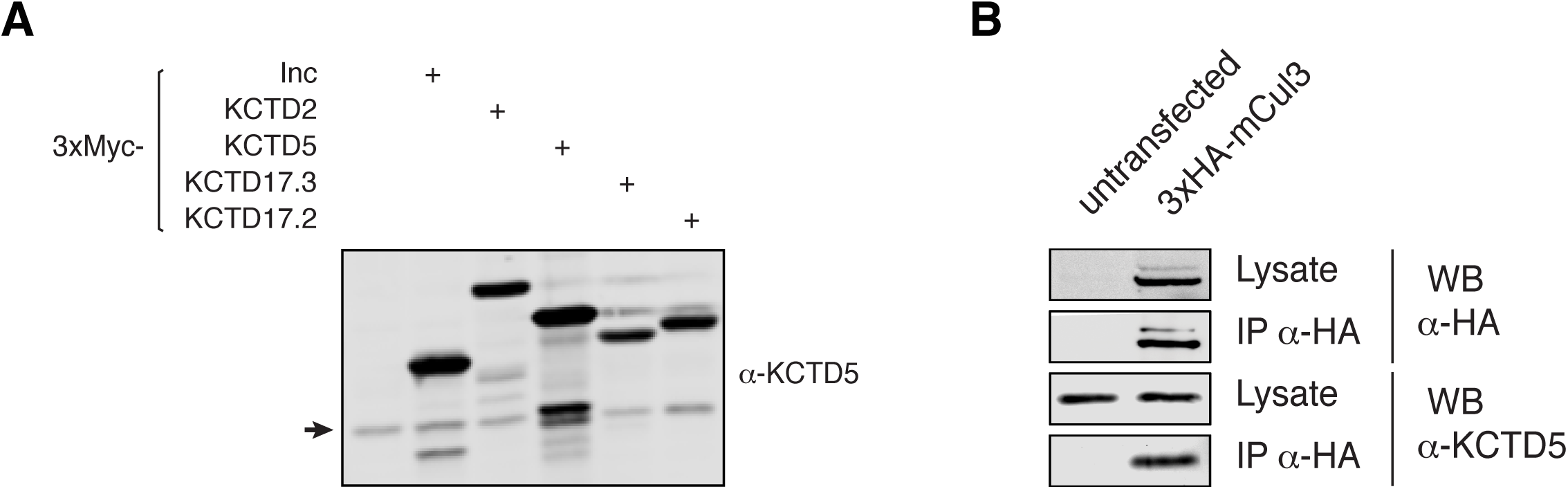
Specificity of anti-KCTD5 antisera and co-immunoprecipitation of Cul3 and endogenous KCTD2/5/17. **(A)** Western blot of extracts from 293T cells transfected with indicated expression vectors and probed with anti-KCTD5. Arrow indicates endogenous species corresponding to KCTD 2/5/17. The first lane of the blot contains a control sample from cells transfected with empty vector and treated in parallel. **(B)** 293T cells transfected and immunoprecipitated as indicated.

**Fig S4.**
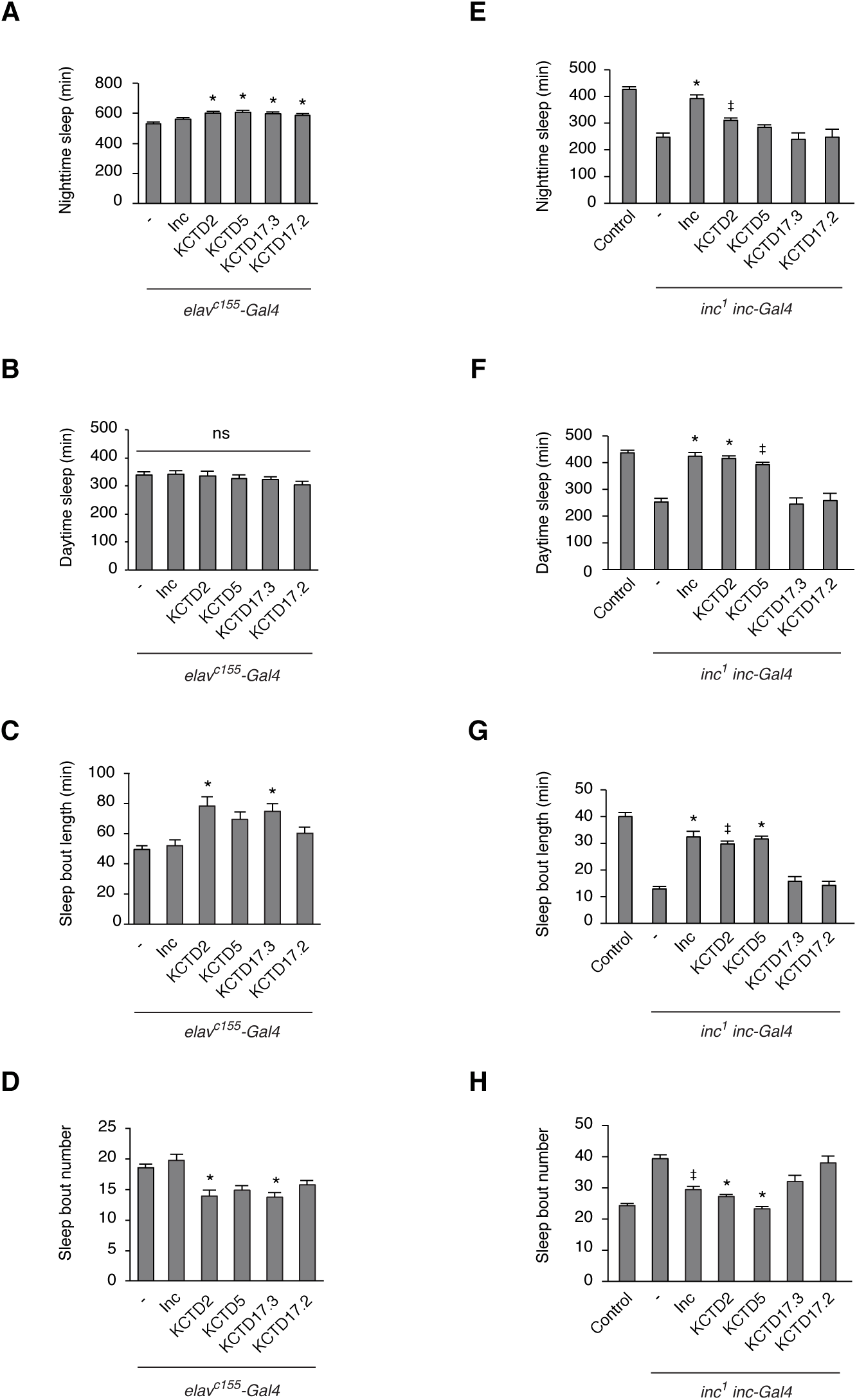
Additional sleep parameters for animals expressing Inc and Inc orthologs. **(A-D)** Sleep parameters for animals expressing Inc and Inc orthologs panneuronally under*elav-Gal4* control. n = 37-40 as in Figure 4C; * p < 0.01 compared to *elav-Gal4* control; ns, not significant (p > 0.05). **(E-H)** Sleep parameters for *inc*^*1*^ *inc-Gal4* animals expressing Inc and Inc orthologs. n=18-157 as in Figure 4D. * p < 0.01 compared to *inc*^*1*^ *inc-Gal4* animals, but not significantly different from wild-type control. ^‡^ p < 0.01 for comparisons to *inc*^*1*^ *inc-Gal4* animals and to wild-type controls. For all panels, mean ± SEM is shown. (A and E) Nighttime sleep. (B and F) Daytime sleep. (C and G) Sleep bout length. (D and H) Sleep bout number.

**S5 Fig.**
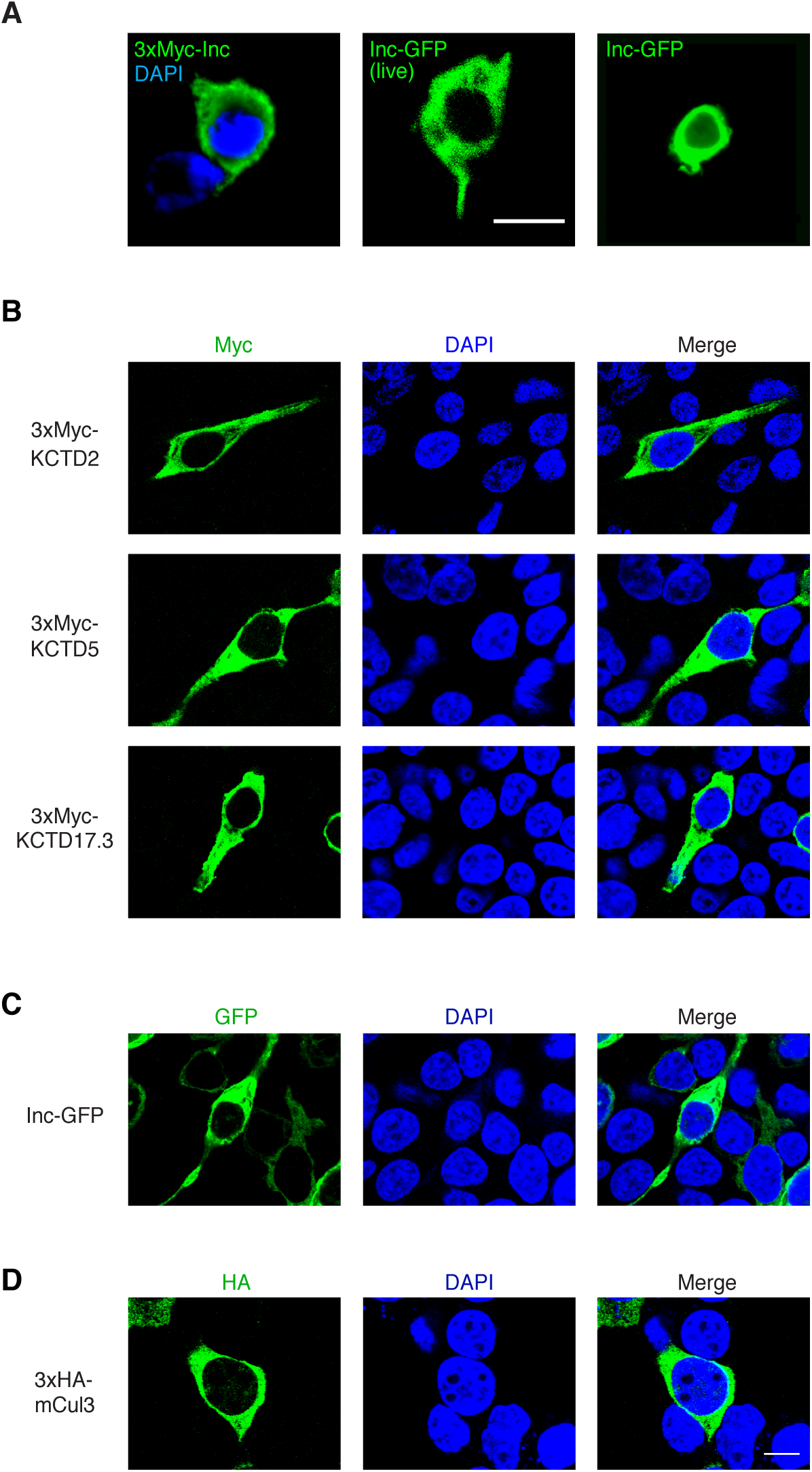
Localization of Insomniac family members in cultured cells. **(A)** Confocal micrographs of fixed S2 cell expressing 3×Myc-Inc (left panel), live S2 cell expressing Inc-GFP (middle panel), and fixed S2 cell expressing Inc-GFP (right panel). **(B-D)** Confocal micrographs of fixed 293T cells expressing indicated proteins bearing a 3×Myc tag (B), GFP tag (C), or 3×HA tag (D). Scale bar is 10 µM.

**S6 Fig.**
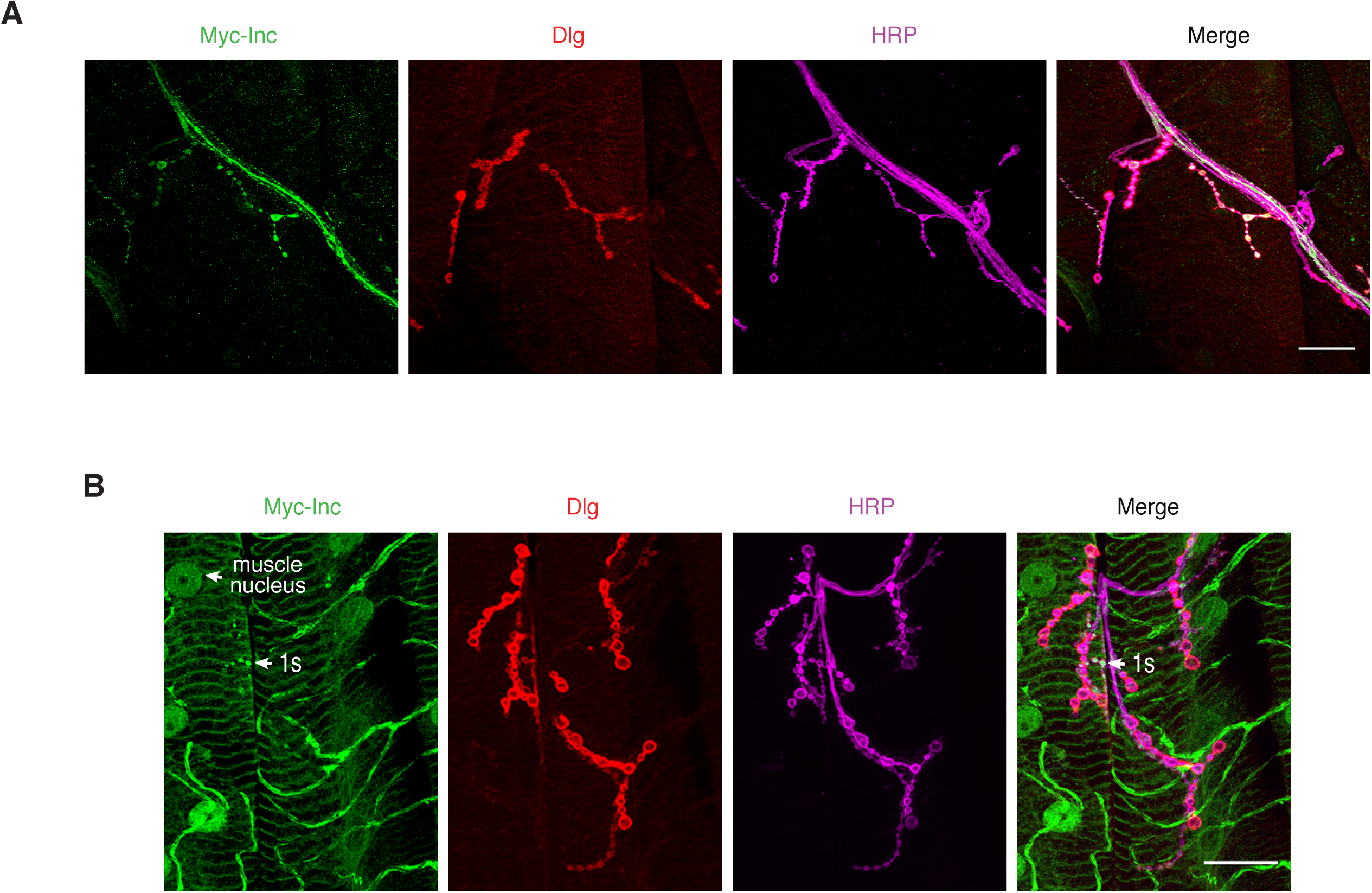
Immunohistochemical analysis of Inc localization at NMJ 4 and in male larvae. **(A)** Confocal micrograph of NMJ 4 from a female third instar *inc-Gal4 / + ; UAS-Myc-Inc / +* larva. (B) Confocal micrograph of NMJ 6/7 from a male third instar *inc-Gal4; UAS-Myc-Inc / +* larva. Note higher level of Inc signal in muscle and HRP-negative trachea relative to female larva shown in Figure 6I; this higher level of expression may reflect dosage compensation of the X-linked *inc-Gal4* transgene in hemizygous males versus heterozygous females, or sex-specific position effects of the X-linked *inc-Gal4* transgene insertion site. Male and female animals heterozygous for autosomal insertions of the *inc-Gal4* transgene exhibit similarly weak levels of *UAS-Myc-Inc* expression in muscle (not shown).

